# Convergent selection on hormone signaling shaped social evolution in bees

**DOI:** 10.1101/2021.04.14.439731

**Authors:** Beryl M. Jones, Benjamin E.R. Rubin, Olga Dudchenko, Callum J. Kingwell, Ian M. Traniello, Z. Yan Wang, Karen M. Kapheim, Eli S. Wyman, Per A. Adastra, Weijie Liu, Lance R. Parsons, S. RaElle Jackson, Katharine Goodwin, Shawn M. Davidson, Matthew J. McBride, Andrew E. Webb, Kennedy S. Omufwoko, Nikki Van Dorp, Mauricio Fernández Otárola, Melanie Pham, Arina D. Omer, David Weisz, Joshua Schraiber, Fernando Villanea, William T. Wcislo, Robert J. Paxton, Brendan G. Hunt, Erez Lieberman Aiden, Sarah D. Kocher

**Author notes:** These authors contributed equally (Beryl M. Jones, Benjamin E.R. Rubin, Olga Dudchenko). These authors contributed equally (Erez Lieberman Aiden, Sarah D. Kocher).

## Abstract

Sweat bees have repeatedly gained and lost eusociality, a transition from individual to group reproduction. Here, we generate chromosome-length genome assemblies for 17 species and identify genomic signatures of evolutionary trade-offs associated with transitions between social and solitary living. Both young genes and regulatory regions show enrichment for these molecular patterns. We also identify loci that show evidence of complementary signals of positive and relaxed selection linked specifically to the convergent gains and losses of eusociality in sweat bees. This includes two proteins that bind and transport juvenile hormone (JH) – a key regulator of insect development and reproduction. We find one of these JH binding proteins is primarily expressed in subperineurial glial cells that form the insect blood-brain barrier and that brain levels of JH vary by sociality. Our findings are consistent with a role of JH in modulating social behavior and suggest eusocial evolution was facilitated by alteration of the proteins that bind and transport JH, revealing how an ancestral, developmental hormone may have been co-opted during one of life’s major transitions. More broadly, our results highlight how trade-offs have structured the molecular basis of eusociality in these bees and demonstrate how both directional selection and release from constraint can shape trait evolution.

---

Organisms situated at the inflection point of life’s major evolutionary transitions provide a powerful framework to examine the factors shaping the evolution of traits associated with these transitions^1,2^. Halictid bees (“sweat bees”, Hymenoptera: Halictidae) offer a unique opportunity to study the evolution of eusociality (social colonies with overlapping generations and non-reproductive workers), since within this group there have been two independent gains^3^ and a dozen losses^4^ of eusociality. As a result, closely related halictid species encompass a broad spectrum of social behavior, from solitary individuals that live and reproduce independently to eusocial nests where individuals cooperate to reproduce as a group, and even polymorphic species that produce both solitary and social nests^5^. This evolutionary replication enables a comparative approach to identify the core factors that shape the emergence and breakdown of eusociality.

To identify these factors, we generated 15 *de novo* genome assemblies and updated 2 additional assemblies in halictids, all with chromosome-length scaffolds (Fig. 1; Fig. S1). We selected species with well-characterized social behaviors that encompass both origins and six repeated losses of eusociality (based on a much broader phylogeny used to reconstruct the origins of eusociality in this group^3^). Sampling closely-related eusocial and solitary species alongside known non-eusocial outgroups provides a powerful framework to examine the molecular mechanisms shaping the evolution of social behavior in these bees^1^.

**Fig. 1.**
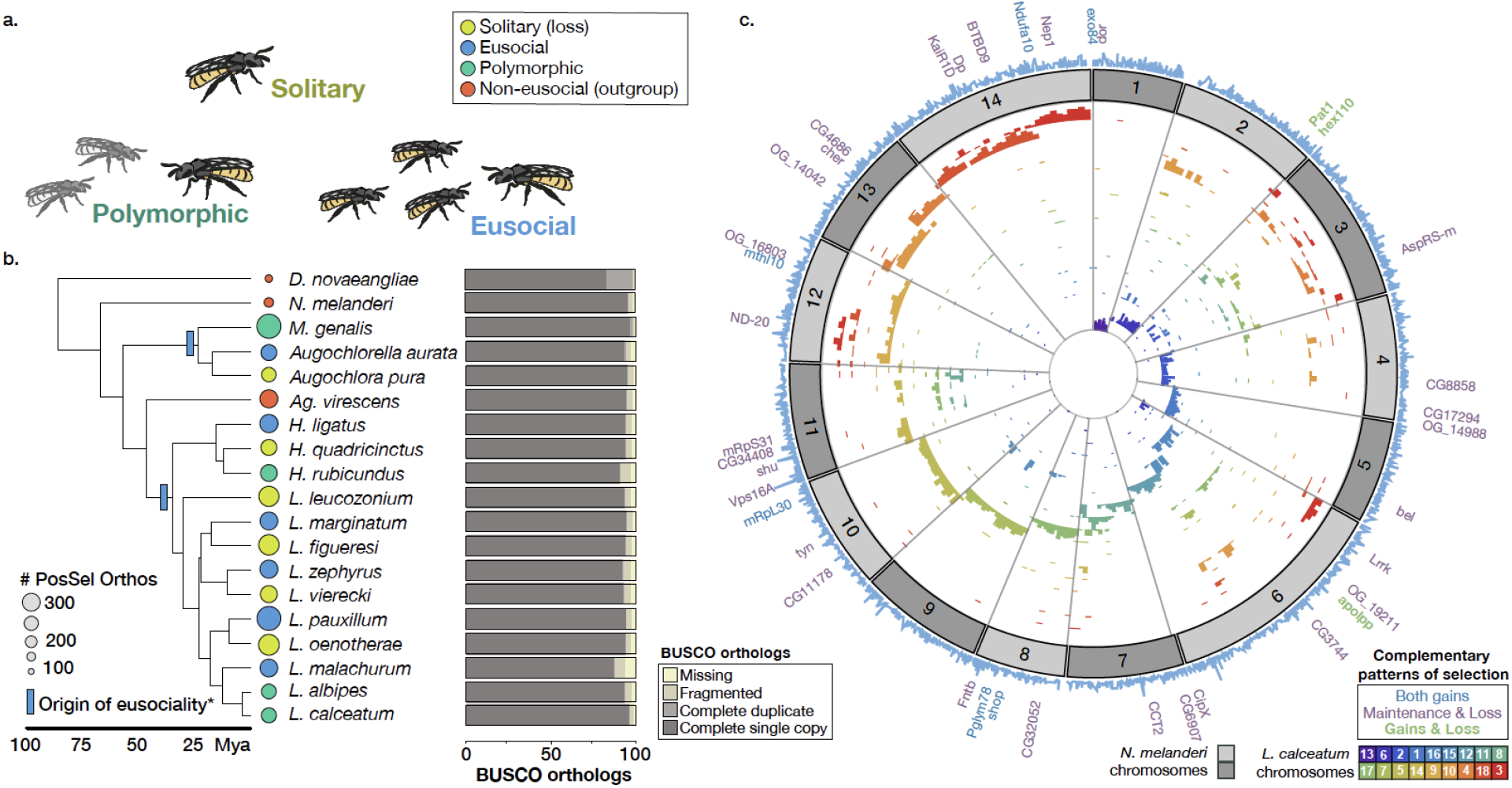
Comparative genomic resources for halictid bees. (**A**) Halictid bees encompass a wide range of behaviors, including solitary (yellow), eusocial (blue) and polymorphic species (green) capable of both solitary and eusocial reproduction. Non-eusocial outgroups (red) reproduce independently. (**B**) Pruned tree showing taxa included in this dataset. Colors at tips indicate species’ behavior, circle sizes proportional to the number of orthologs under positive selection on each terminal branch (abSREL, HyPhy^18^ (FDR<0.05). Light blue rectangles denote gains of eusociality (*inferred from much broader phylogenies than shown here^3^). Proportions of complete/fragmented/missing BUSCO orthologs are shown for each genome. (**C**) Genomic data aligned to *Nomia melanderi* (NMEL). The 14 NMEL chromosomes are represented as a circular ideogram with consecutive chromosomes shown in alternating dark/light gray. The inner spiral comprises 18 color-coded tracks, each corresponding to one *L. calceatum* (LCAL) chromosome; the y-axis represents the frequency of regions aligning to the corresponding region in NMEL. Most alignments fall into a single “wedge”, indicating that each LCAL chromosome corresponds to just one NMEL chromosome, a pattern typical for sweat bees and unlike that of mammals (Fig. S2-S3). Outer blue line plot indicates number of branches where positive selection was detected at each gene (abSREL, FDR<0.05), and gene names shown are those with convergent/complementary patterns of selection: positive selection at both origins (blue), intensification of selection in extant eusocial lineages and relaxation of selection in secondarily solitary species (HyPhy RELAX, FDR<0.1; purple), and complementary patterns of both positive selection on the origin branches and convergent relaxation of selection with losses of eusociality (green). Not all genes in these categories are annotated in NMEL and are therefore not labeled.

We searched for signatures of positive selection associated with the convergent gains of eusociality as well as signatures of relaxed selection when eusociality is lost. These complementary patterns indicate genomic loci that are associated with costs or trade-offs underlying the maintenance of social traits. We find that some of the targets of selection implicated in the origins and elaborations of eusociality, such as young, taxonomically restricted genes^6^ and gene regulatory elements^7^, also show relaxation of selective pressures when social behavior is lost. In addition, we uncovered four genes strongly associated with the evolution of eusociality in halictid bees, including the two primary juvenile hormone binding proteins (JHBPs): *apolipoprotein*^8–10^ and *hexamerin110*^11,12^. Using single-cell RNA-sequencing, we localized the expression of *apolipoprotein* in the brain to glial cells involved in forming the insect “blood-brain barrier”^13^. In addition, we find evidence that eusocial reproductive females have increased levels of JH III in their brains compared to their solitary counterparts; this could potentially be mediated by changes in the transport of JH. These results provide new insights into how JH signaling may have been modified to shape the evolution of eusociality.

## Results

We generated new comparative genomic resources for studying the evolution of eusociality in halictid bees. Genome assemblies of 17 species ranged in size from 267 to 470 Mb, with estimated numbers of chromosomes ranging from 9 to 38 (Table S1). Broadly, we found that, in contrast to mammalian species (Fig. S2), genomic rearrangements among the bee species occur disproportionately within rather than between chromosomes (see Fig. 1c, Fig. S3). Consequently, loci that are on the same chromosome in one bee species also tend to occur on the same chromosome in other bee species. This observation is similar to previous findings in dipteran genomes^14,15^ and may indicate a broader trend during the evolution of insect chromosomes. To increase the quality of genome annotations, we generated tissue-specific transcriptomes for 11 species, and we also sequenced and characterized 1269 microRNAs (miRs) expressed in the brains of 15 species (Table S2). The number of annotated genes ranged from 11,060 to 14,982, and BUSCO^16^ analyses estimated the completeness of our annotations to range from 93.4 to 99.1% (Fig. 1b; Table S1). Whole-genome alignments were generated with progressiveCactus^17^ (Fig. 1c), which we then used to identify 52,421 conserved, non-exonic elements present in 10 or more species. All genomes, annotations, and alignments can be viewed and downloaded from the Halictid Genome Browser (https://beenomes.princeton.edu).

### Young genes and gene regulatory expansions are associated with trade-offs underlying the maintenance of eusociality

Previous studies of eusociality have suggested that, similar to their importance in the evolution of other novel traits^19^, younger or taxonomically-restricted genes (TRGs) may play key roles in the evolution of eusocial behavior^20^. To test this hypothesis, we examined the relationship between gene age and selection associated with eusocial origins, maintenance, and reversions to solitary life histories. We found a greater proportion of young genes compared with old genes experience relaxed selection when eusociality is subsequently lost (Fig. 2a; Pearson’s r=0.869, p=0.002). This relationship does not hold for orthologs showing evidence of relaxed selection on eusocial branches, nor are younger genes more likely than older genes to experience intensified selection associated with either the origins or maintenance of eusociality (Fig. S4; Table S3).

**Figure 2.**
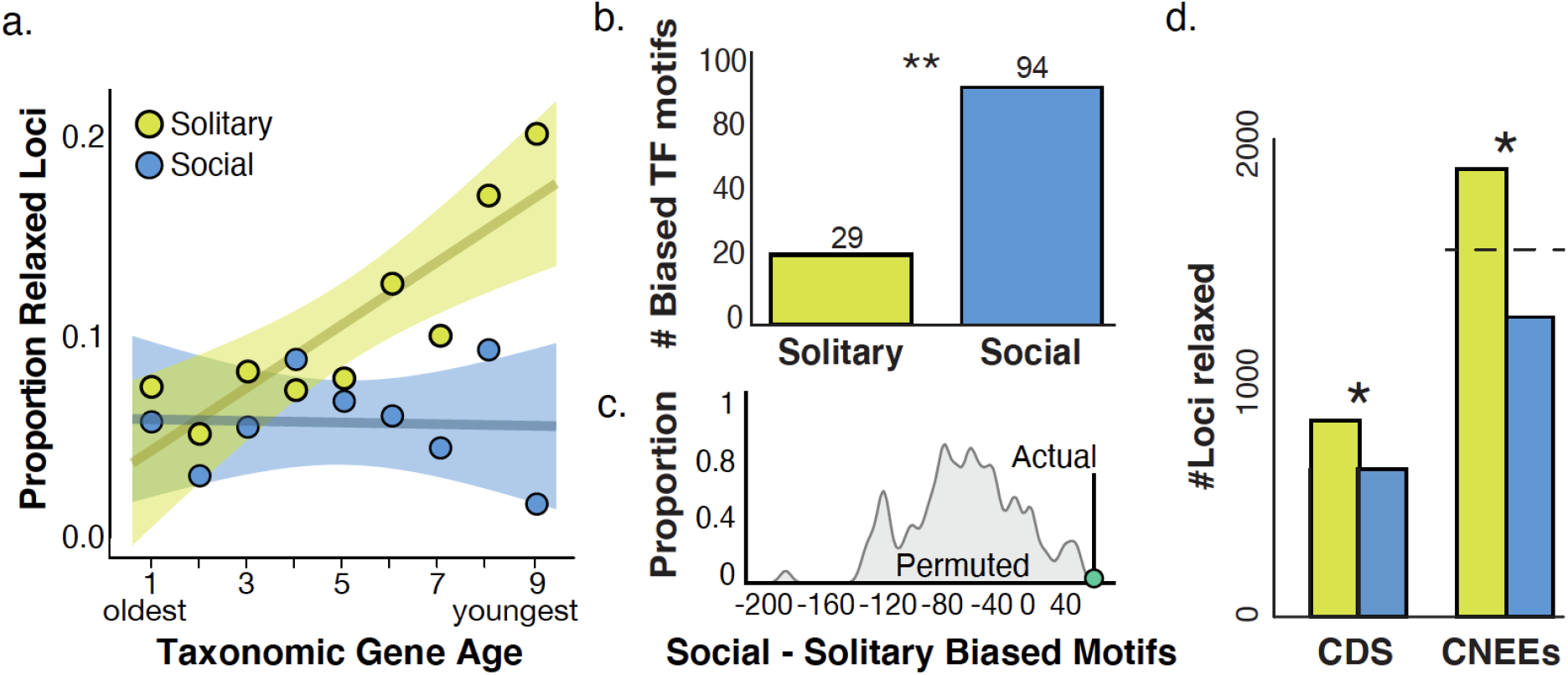
The maintenance of eusociality is associated with young genes and gene regulation. (**A**) Younger genes are more likely to show signatures of relaxed selection when social behavior is lost. Circles show the proportion of genes in each age class that show evidence of relaxed selection in solitary (yellow) or social (blue) lineages, from the oldest Bilaterian group (Age=1) to the youngest, halictid-specific taxonomically restricted genes (Age=9). For solitary lineages, gene age is significantly correlated with the proportion of those genes with evidence of relaxed selection (Pearson correlation, r=0.869, p=0.002; social lineages: r=-0.0447, p=0.91). Shading represents 95% confidence intervals. (**B**) Stubb scores^22^ were calculated for 223 Drosophila transcription factor binding motifs in each genome, and each motif was tested for overrepresentation in solitary/social genomes; 94 motifs were enriched in social genomes compared to 29 enriched in solitary genomes. “**” indicates p<0.01. (**C**) Permutation tests reveal the ∼3-fold enrichment in (B) is unlikely to occur by chance (empirical p<0.01). (**D**) Taxa that have lost eusociality have higher proportions of loci experiencing relaxed selection after phylogenetic correction, both for coding sequences (CDS; Fisher’s exact, p=2.42×10-7, odds-ratio=1.48) and for conserved, non-exonic elements (CNEEs). Social lineages have fewer fast-evolving CNEEs than chance (Binomial test, p<1×10-10), while solitary taxa have more than chance (Binomial test, p=5.27×10-9). The dashed line indicates the null expectation for CNEEs, and * indicates significant differences in the number of loci between eusocial and solitary species.

Gene regulatory changes have also been implicated in eusocial evolution^21^, including the expansion of transcription factor (TF) motifs in the genomes of eusocial species compared with distantly-related solitary species^7^. To assess the degree to which changes in gene regulation may facilitate the evolution of social behavior in halictids, we characterized TF motifs in putative promoter regions in each halictid genome. For each species, we defined these regions as 5kb upstream and 2kb downstream of the transcription start site for each gene^7^ and calculated a score for a given TF in each region that reflects the number and/or strength of predicted binding sites^22^.

If social species have a greater capacity for gene regulation compared to lineages that have reverted to solitary nesting, then we would expect to find more motifs with scores (reflecting both strength and number of binding sites) that are higher in social taxa compared to secondarily solitary taxa. In support of this hypothesis, we find a greater than 3-fold enrichment of TF motifs that are positively correlated with social lineages compared to secondarily solitary lineages after phylogenetic correction (Fig. 2b; Table S4), and permutation tests indicate that this difference is highly significant (Fig. 2c). Five of these socially-biased motifs were previously identified as associated with eusocial evolution in bees^7^, including the motifs for *lola, hairy, CrebA, CG5180*, and the *met/tai* complex, which initiates downstream transcriptional responses to JH. Moreover, conserved, non-exonic elements (CNEEs) showed a bias toward faster rates of evolution in secondarily solitary species compared with eusocial lineages (Fig. 2d). This acceleration is likely to be a signature of relaxed constraint in solitary lineages, further supporting the role of gene regulation in the maintenance of eusocial traits.

### Convergent and complementary signatures of selection are associated with the gains and losses of eusociality

To determine if specific loci show convergent signatures of selection associated with the evolution of eusociality, we identified genes with positive selection on each branch representing a gain of eusociality. We found 309 genes on the Augochlorini origin branch and 62 genes on the Halictini origin branch with evidence of position selection (Fig. 3; Tables S5). On the Augochlorini branch, genes with signatures of positive selection were enriched for cell adhesion (GO:0007155, Fisher’s Exact, q=2.01e-4; enrichment=2.92; Table S6). This list included *taiman*, which encodes a protein that co-activates the bHLH-PS transcription factor and JH receptor, Met^23–27^ (Table S5). There was no detectable gene ontology (GO) enrichment among positively-selected genes on the Halictini branch (Fisher’s Exact, q>0.1). Nine genes showed signatures of positive selection on both branches (Table S5). These genes include *shopper*, which is involved in neuronal network function and the ensheathing of glial cells^28^, and *ND-42*, a suppressor of Pink1 which is associated with neurocognitive functioning^29^.

**Figure 3.**
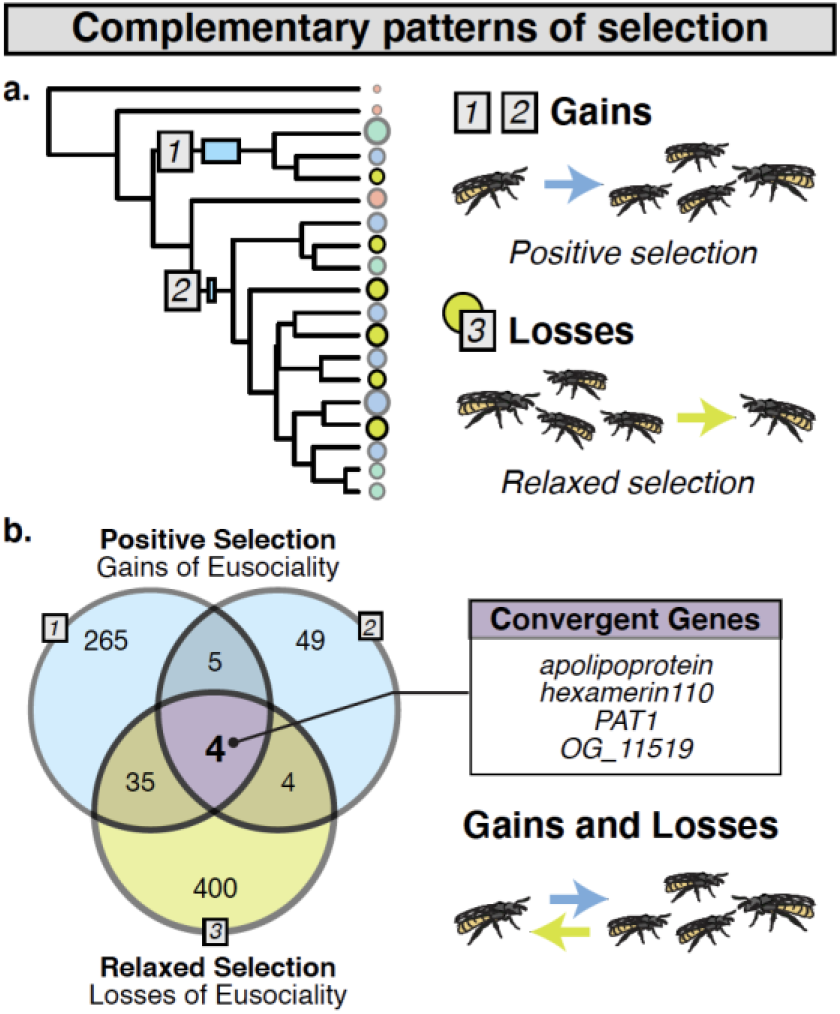
Signatures of selection associated with the gains and losses of eusociality in halictids. (**A**) Orthologs were tested for evidence that dN/dS >1 at a proportion of sites on focal branches (Gains 1 and 2 denoted with blue squares 1-2; abSREL^18^ tests in HyPhy, FDR<0.05) and for evidence of relaxed selection (gray square 3, yellow circles; HyPhy RELAX^33^, FDR<0.1). (**B**) Nine loci overlapped both origins (enrichment ratio=3.97, Fisher’s exact p=0.004). Four orthologs overlapped all 3 tests (Multi-set Exact Test^36^; fold-enrichment=8.85, p=0.001).

A unique attribute of halictid bees is that there have been a number of independent losses of eusociality^30^ in addition to repeated gains^3^. These reversals provide a powerful lens to identify key genomic factors needed for the maintenance of social living because organisms are expected to minimize investment in traits when social behaviors are lost or unexpressed. This results in the reduction or removal of selective pressures previously maintaining these costly but essential traits^31,32^. Thus, we predicted that genes associated with trade-offs or costly roles in maintaining eusocial societies should show relaxation of constraint in species that have secondarily reverted to solitary nesting. Consistent with this hypothesis, we found 443 genes showing evidence of convergent, relaxed selection on the six branches representing independent losses of eusociality (HyPhy RELAX^33^, FDR<0.1; Table S5). These genes are enriched for chromosome condensation (GO:0030261, Fisher’s Exact, q=0.067, enrichment=4.09), indicating that they may play a role in chromosome accessibility and gene regulation^34^. They are also enriched for vacuolar transport (GO:0007034, Fisher’s Exact, q=0.067, enrichment=3.11; Table S6).

To determine whether or not this pattern is unique to the loss of eusociality, we ran the same tests for relaxed selection using extant eusocial lineages as the focal branches. We found 305 genes with signatures of relaxation in eusocial species (HyPhy RELAX^33^, FDR<0.1; Table S5) enriched for four GO terms related to metabolism (Table S6). This is a significantly lower proportion of genes experiencing relaxed selection in eusocial species compared to those experiencing relaxed selection among solitary species (Fisher’s exact test p=2.42×10^−7^, odds-ratio=1.48), suggesting that the loss of eusociality is more often associated with a release of constraint compared with eusocial maintenance or elaboration.

We also identified 34 genes that show intensification of selection on extant eusocial lineages and relaxation in secondarily solitary species (HyPhy RELAX, FDR<0.1 for both tests). The convergent intensification of selection on eusocial lineages suggests that these genes are likely to be particularly relevant to the maintenance or elaboration of eusociality. They are enriched for regulation of SNARE complex assembly (GO:0035542, Fisher’s Exact, q=0.074, enrichment=80.91; Table S5), which is a key component of synaptic transmission that has also been implicated with variation in social behavior in *L. albipes*^2^ and wasps^35^.

By comparing genes associated with the emergence of eusociality to those associated with its loss, we have the unique ability to identify some of the most consequential molecular mechanisms shaping social evolution. If a shared set of genes is associated with the emergence and maintenance of social behavior in this group, then we would expect to find genes experiencing both positive selection when eusociality emerges and relaxed selection when social behavior is lost (Fig. 3b). Indeed, we find four genes matching these criteria: *OG_11519*, a gene with no known homologs outside of Hymenoptera, *Protein interacting with APP tail-1* (*PAT1), apolipoprotein* (*apolpp*), and *hexamerin110* (*hex110*; Fig. 3b). This overlap is significantly more than expected by chance (Multi-set Exact Test^36^; fold-enrichment=8.85, p=0.001). *OG_11519* has no identifiable protein domains but is conserved throughout the Hymenoptera. PAT1 modulates mRNA transport and metabolism^37^, and ApoLpp and Hex110 have been established as the primary JH binding proteins (JHBPs) across multiple insect orders^9–11,38^. These complementary patterns of positive selection when eusociality arises and relaxation of selection when it is lost suggests that this small but robust set of genes is associated with costly trade-offs linked to the evolution of eusociality^32^.

### Genes that mediate juvenile hormone binding and transport show evidence of convergent and complementary selection

Two of the 4 genes that show signatures of both positive selection when eusociality is gained and relaxed selection when eusociality is lost (*apolpp* and *hex110*) encode the primary binding proteins for JH, an arthropod-specific hormone with pleiotropic effects on numerous insect life history traits^39^. To further investigate the evidence for selection on these genes, we implemented a mixed-effects maximum likelihood approach (MEME^40^) and found region-specific, faster rates of evolution on eusocial branches compared to non-eusocial outgroups for both proteins. Sites with evidence of positive selection are present in the functional regions of both proteins (Fig. S5), including the receptor binding domain and predicted binding pocket for ApoLpp as well as in all three Hemocyanin domains of Hex110 (associated with storage functions^41^) and its predicted binding pocket. Though both of these proteins are highly pleiotropic^41,42^, their shared role in JH binding and transport suggests that positive selection may have shaped JHBP function as eusociality emerged in two independent lineages of halictids and that some of these changes may also be associated with costs when eusociality is lost.

### Characterization of JHBP expression patterns and JH III levels in the brain

While associations between circulating JH levels and division of labor are well established in the social insects^43^, we still do not understand which components of JH signaling pathways have been targeted by natural selection to decouple JH from its ancestral role in development and reproductive maturation^44^ and generate new links between JH and social traits. Because ApoLpp delivers cargo to target tissues and has been shown to cross the blood-brain barrier^45^, we hypothesized that differences in ApoLpp transport and the availability of JH in the brain can help generate novel relationships between JH and behavior. To examine the potential role of cell type-specific expression of ApoLpp in modulating JH signaling in the brain, we generated a single-cell RNA-Sequencing (scRNA-Seq) brain dataset using two sweat bee species: *L. albipes* and *L. zephyrus* (Fig. 4a). We identified one cluster, characterized by markers of glial cells, to be the primary location of *apolpp* brain expression (Fig. 4b). In addition, three related lipid transfer-associated genes, *apolipoprotein lipid transfer particle* (*apoLTP*), *megalin* (*mgl;* experiencing relaxed selection in solitary lineages; Table S5), and *Lipophorin receptor 1 (LpR1*), as well the JH degradation enzyme *jheh2*, are expressed primarily in these glial cells (Fig. 4b). Further subclustering demonstrates that both *apolpp* and *apoLTP* are primarily expressed in a subperineurial glia-like cluster (Fig. 4b); subperineurial glia, along with perineurial glia, form and regulate the permeability of the blood brain barrier in *D. melanogaster*^46,47^.

**Figure 4.**
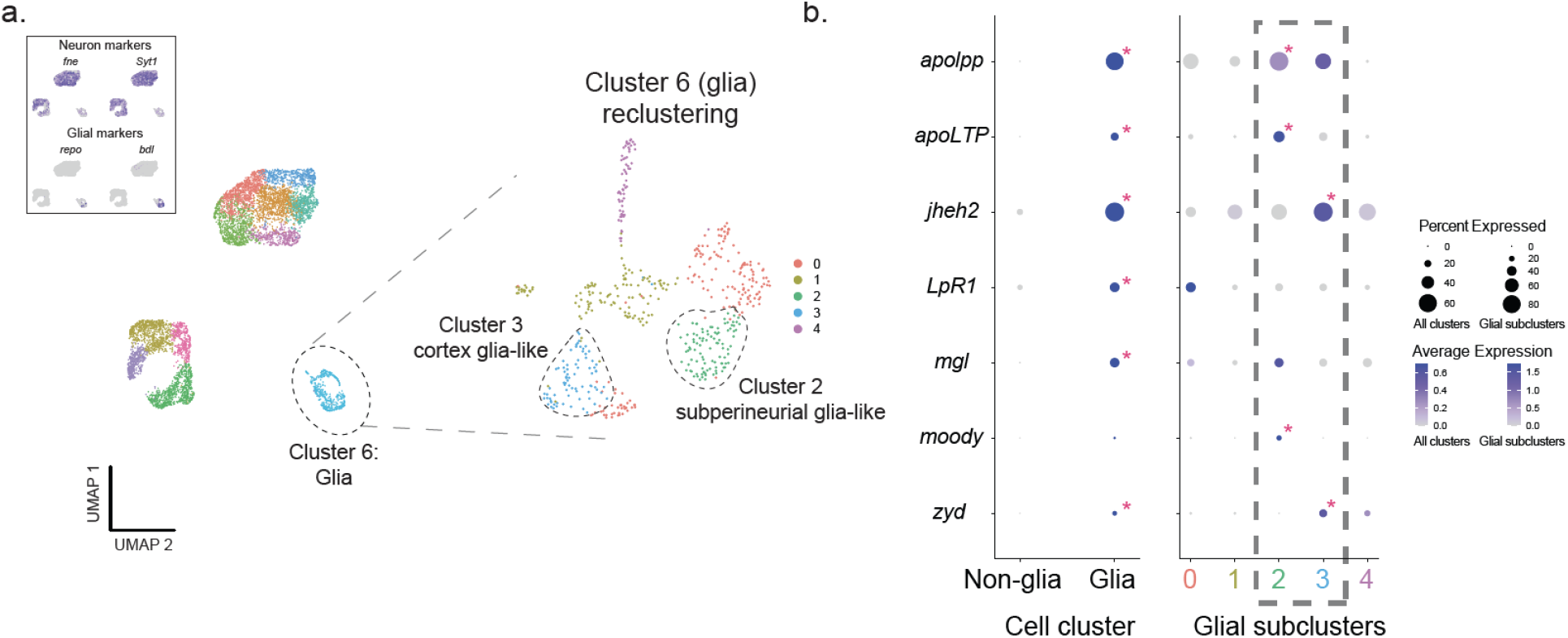
*apolpp* and associated lipid transport genes are expressed in glial cells. (A) Single-cell RNA sequencing (scRNAseq) of the halictid brain. Clustering and visualization of ∼7,000 cells from 4 halictid brain samples revealed 11 cell clusters. Expression of canonical markers of insect neurons and glia showed clear segregation of glia into a single cluster, Cluster 6. (B) Four genes associated with lipid binding, including *apolpp, apoLTP, LpR1, and mgl*, as well as *jheh2*, were co-expressed in this cluster. Focal subclustering of Cluster 6 revealed that *apolpp* and *Apoltp* are co-expressed with *moody*, a marker of subperineurial glia (Cluster 2), and *jheh2* is co-expressed with *zyd*, a marker of cortex glia (Cluster 3). Pink asterisks in dotplots indicate that a specific gene (row) is significantly upregulated in a given cell cluster compared to others (column). All expression values represent gene counts following sequencing depth normalization.

Our selection results suggest that JH transport to the brain may play a critical role in eusocial evolution in halictids. We used liquid chromatography – mass spectrometry (LC-MS) to measure JH III titers in dissected brains and found higher concentrations of JH III in social (*A. aurata*) foundresses compared with solitary (*A. pura*) foundresses; social *A. aurata* workers appear to have intermediate levels of JH III (Fig. 5a). Next, we used topical, abdominal applications of isotopically labeled JH III-d3 to demonstrate that the bee brain is permeable to JH III (Fig. 5b; Fig. S6). Taken together, our results suggest differential responses to JH in the brain and other tissues could be associated with behavioral polyphenisms among the social insects^44^.

**Figure 5.**
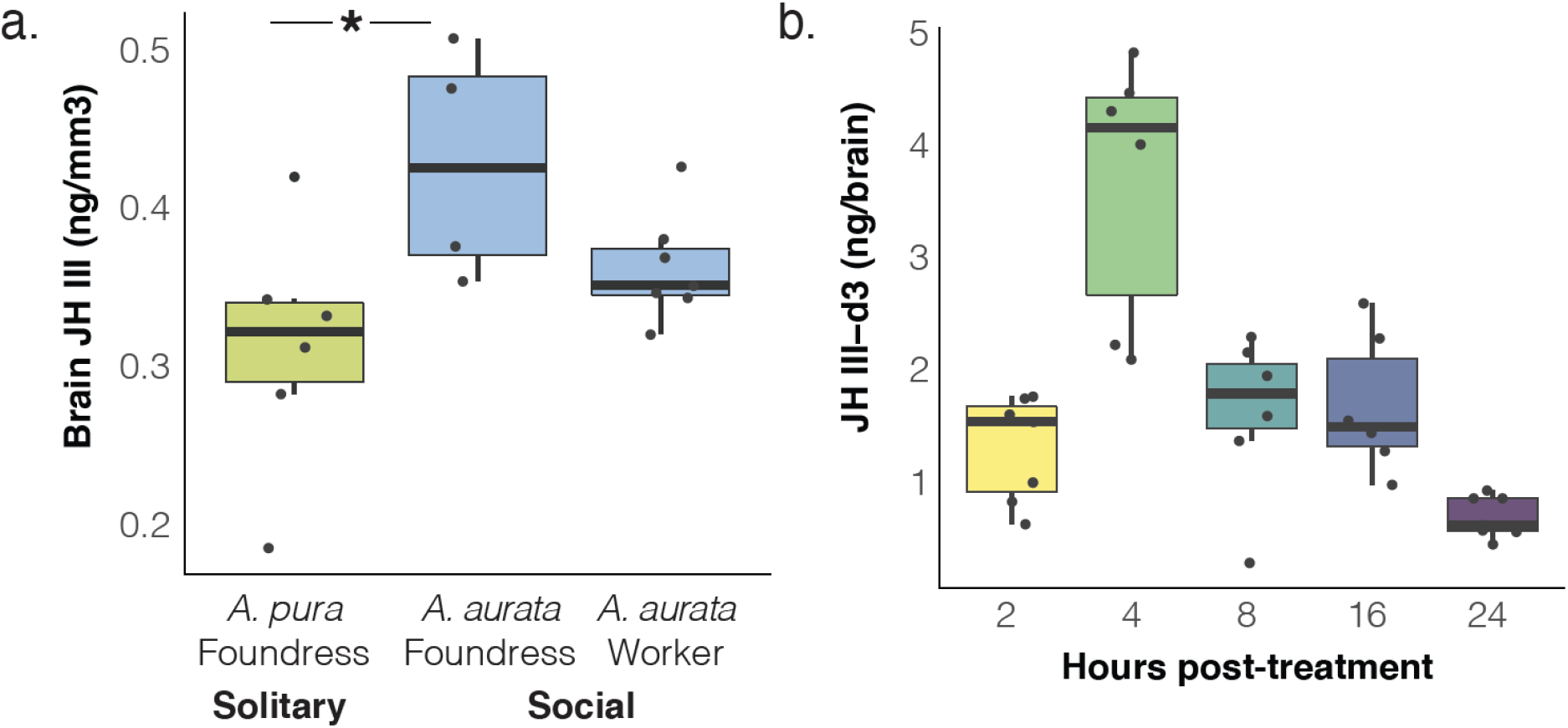
JH is higher in brains of eusocial foundresses and crosses the blood-brain barrier. (**A**) LC/MS quantification of JH III in the brains of solitary *A. pura* (n=6) and eusocial *A. aurata* foundresses (n=4) and workers (n=7) reveal higher endogenous levels of JH III in social foundresses (Wilcoxon rank sums test, p=0.029; *asterisks indicate post-hoc, pairwise comparisons, p<0.05). (**B**) Isotopically-labeled JH III-d3 applied to the abdomens of *A. aurata* is detected in the brain as soon as 2h later and peaks at 4 hours post-treatment (n=7 for 2,8,24h; n=6 for 4h and 16h).

## Discussion

We leveraged the powerful natural variation in social behavior of sweat bees and developed new comparative genomic resources to identify mechanisms shaping transitions in social evolution. In addition to multiple gains of eusociality, halictids provide an excellent opportunity to study genomic signatures of eusocial loss, with repeated, recent reversions from eusocial to solitary life history strategies. By studying both gains and losses within this group, we have uncovered multiple genome-wide patterns as well as specific targets of selection associated with eusociality, a major evolutionary transition.

First, we tested for broad patterns in genome evolution that have previously been implicated in the elaboration or maintenance of eusociality, including a role for younger or taxonomically-restricted genes (TRGs)^20^ and changes in gene regulation^7,21^. We found several lines of evidence suggesting these patterns extend to social evolution in sweat bees. Our finding that TRGs are disproportionately experiencing a relaxation of selection pressure when social behavior is lost suggests that, in addition to influencing eusocial maintenance, younger genes are associated with costs or trade-offs linked to eusociality. These findings support studies of complex eusocial hymenopterans, including honey bees, ants, and wasps, in which young TRGs have been linked to the evolution of non-reproductive workers^20^. Our work extends this body of evidence to suggest that evolutionary changes in these TRGs may also incur costs when lineages revert to solitary strategies. We also find evidence that, like what has been found in other social lineages^7,21,48^, changes in gene regulation are associated with the elaboration of eusocial traits. TF motifs are expanded in social halictid genomes compared with secondarily solitary lineages, implicating more complex gene regulatory networks associated with eusociality. Further, many putative regulatory regions of halictid genomes (CNEEs) show faster rates of evolution in secondarily solitary lineages. This acceleration is likely to be a signature of relaxed constraint in solitary lineages, further supporting the role of gene regulation in the maintenance of eusocial traits.

In addition to the genome-wide patterns associated with eusocial transitions, the repeated gains and losses of eusociality among halictids enabled us to probe the most consequential molecular mechanisms associated with social evolution. We identified a small but robust set of genes with complementary signatures of selection linked to these social transitions. These complementary patterns highlight the importance of these genes in eusocial lineages: convergent signatures of relaxed selection when eusociality is lost suggest they are also associated with costly trade-offs^32^. Strikingly, 2 of these 4 genes (*apolpp* and *hex110*) encode the primary binding proteins for juvenile hormone (JH), a hormone that regulates many aspects of insect life history including development, reproduction, diapause and polyphenism^43,49^. Together, *apolpp* and *hex110* are thought to bind nearly all JH in insect hemolymph^38^. We identified positive selection on sites present in the functional regions of both ApoLpp and Hex110, with faster rates of evolution on eusocial branches compared with non-eusocial outgroups.

Our single-cell transcriptomics dataset reveals that *apolpp* is primarily expressed in the brain in a cell cluster that co-expresses markers of glial cells, three related lipid transfer-associated genes, and the JH degradation enzyme, *jheh2*. In *Drosophila*, ApoLpp forms a complex with other lipoprotein particles and can cross the blood-brain barrier^45^. In our halictid dataset, *apolpp* was enriched in the cell cluster expressing markers of subperineurial glia, a glial subtype that contributes to the formation and permeability of the blood-brain barrier, suggesting ApoLpp may have a similar role in mediating transport of cargo to the brain in bees. Though JH has previously been quantified in whole-insect heads^50^, it has not yet been quantified directly in the insect brain. We used isotopically labeled JH to demonstrate that JH is able to cross the blood-brain barrier in bees, including halictids. In addition, we found that endogenous levels of JH III are higher in the brains of social compared with closely-related solitary female foundresses. Taken together, our results suggest a model where glial expression of ApoLpp and other lipid transport proteins may work in concert with JH degradation enzymes to modulate the uptake and availability of JH to the brain in a way that differentiates behavioral castes of social halictids. Similar hormonal gatekeeping mechanisms have also been recently proposed in ants (Ju & Glastad et al, *unpub*.), suggesting that the regulation of JH in the brain may be a convergently evolved feature of caste differentiation in the social insects.

JH is essential to reproductive maturation in solitary insects, but this signaling system has also been frequently co-opted during major life history transitions^39,49,51,52^, including eusociality^43^. In relatively small eusocial societies like halictids^53,54^, paper wasps^55^, and bumblebees^56,57^, JH has maintained its association with reproductive maturation, but has also gained a new role in mediating aggression and dominance. In the much more elaborate eusocial colonies of honey bees, further modifications to JH signaling have also resulted in a secondary decoupling of JH in workers independent of its ancestral, reproductive role^43^. Although the association between JH titers and division of labor is well established in these taxa, we still do not understand which components of the JH signaling pathways have been targeted or modified by natural selection to evolutionarily link JH and social traits. Our discovery that JHBPs experience complementary selective pressures when eusociality is gained or lost suggests a mechanism for how this novel association may arise. Specifically, alteration of JHBPs may lead to changes in JH availability in a tissue-specific manner^56^, facilitating the decoupling of JH from its ancestral role in reproduction^44^. Through modification of binding affinity and/or altered cellular uptake of JH, evolutionary changes to *apolpp* and *hex110* could modify the overall and tissue-specific levels of JH^58^, with differential responses to JH leading to behavioral polyphenisms among individuals^44^. Following transport to relevant tissues, JH binds to the JH receptor, Met, which coupled to a co-receptor, Taiman, initiates downstream effects^24^. We found evidence for positive selection on *taiman* linked to the origin of eusociality in the Augochlorini as well as expansion of the *met/tai* TF motif in eusocial lineages. This suggests that modifications to JH response-elements have also helped to fine-tune JH signaling in this group of bees. Future studies are needed to help elucidate the relative influences of modulating JH availability versus refining downstream responses to JH in the origins and elaboration of social traits.

Our findings suggest that modifications to JHBPs along with changes to downstream responses to JH could generate novel relationships between behavioral and reproductive traits – a key feature of the origins and evolution of eusociality^44^. This hypothesis is consistent with theory suggesting that conditional expression of a trait (i.e. eusociality) leads to independent selection pressures and evolutionary divergence^59^; evolution of JHBPs may be one example of such divergence associated with conditional expression of social behavior.

## Conclusions

Sweat bees repeatedly traversed an evolutionary inflection point between a solitary lifestyle and a caste-based eusocial one with multiple gains and losses of this trait. We developed new comparative genomic resources for this group and identified complementary signatures of convergent selection associated with the emergence and breakdown of eusociality. Factors associated with the origins or maintenance of eusociality are also associated with its loss, indicating that there may be trade-offs, constraints, or costs associated with these genomic changes. Strikingly, we find that the functional domains of JHBPs show convergent and complementary signatures of selection as eusociality has been gained and lost in halictids, and that these modifications could affect the transport and availability of JH. Coupled with our finding that JH is present in the insect brain, our results help to explain how novel linkages between social behaviors and endocrine signaling could convergently shape the evolution of eusociality.

## Acknowledgments

We would like to thank our many colleagues that contributed samples and field support to this dataset, including: Jason Gibbs, Sam Droege, Raphael Jeanson, Joan Milam, Jakub Straka, Marion Podolak, James Cane, and Mallory Hagadorn. We also thank Miriam Richards, Cecile Plateaux-Quenu, Jason Gibbs, and Laurence Packer for discussion and insights on halictid life history and behavior. Tim Sackton, Russ Corbett-Detig, Nathan Clark, Adam Siepel, and Xander Xue provided discussion and advice on data analysis. We also thank Wenfei Tong for the bee drawings and Michael Sheehan for assistance with RNA library preparation. This work was supported by NSF-DEB1754476 awarded to SDK and BGH, NIH 1DP2GM137424-01 to SDK, USDA NIFA postdoctoral fellowship 2018-67012-28085 to BERR, DFG PA632/9 to RJP, a Smithsonian Global Genome Initiative award GGI-Peer-2016-100 to WTW and CK, a Smithsonian Institution Competitive Grants Program for Biogenomics (WTW, KMK, BMJ), a Smithsonian Tropical Research Institute fellowship (to CK), and a gift from Jennifer and Greg Johnson to WTW. MFO was supported by Vicerrectoría de Investigación, UCR, project B7287. ELA was supported by an NSF Physics Frontiers Center Award (PHY1427654), the Welch Foundation (Q-1866), a USDA Agriculture and Food Research Initiative Grant (2017-05741), and an NIH Encyclopedia of DNA Elements Mapping Center Award (UM1HG009375). Sampling permit details: SDK, ESW, and MFO (R-055-2017-OT-CONAGEBIO), SDK (P526P-15-04026), and RJP (Belfast City Council, Parks & Leisure Dept).

## Funding

NSF-DEB1754476 (SDK, BGH)

NIH 1DP2GM137424-01 (SDK)

USDA NIFA fellowship 2018-67012-28085 (BERR)

DFG PA632/9 (RJP)

Smithsonian Global Genome Initiative GGI-Peer-2016-100 (WTW, CK)

Smithsonian Institution Competitive Grants Program for Biogenomics (WTW, KMK, BMJ)

STRI fellowship (CK)

A gift from Jennifer and Greg Johnson (WTW)

Vicerrectoría de Investigación UCR B7287 (MFO)

NSF Physics Frontiers Center (PHY1427654) (ELA)

Welch Foundation (Q-1866) (ELA)

USDA Agriculture and Food Research Initiative (2017-05741) (ELA)

NIH Encyclopedia of DNA Elements Mapping Center (UM1HG009375) (ELA)

## Author contributions

Study design: BMJ, BERR, SDK

Initial draft: BMJ, BERR, SDK

Sample collections: BMJ, BERR, ESW, BMJ, SDK, MFO, KMK, RJP, WTW, CJK, KSO

Genomic library generation: BERR, CJK, ZYW, NVD

Hi-C: OD, BGSH, MP, ADO, DW

Genome annotation/alignment: BERR, LP, AEW

Genomic analyses: BMJ, BERR, SDK, OD, KMK, BGH, JS, FV, IMT

Database: AEW

Laboratory experiments: WL, ESW, SRJ, KG, SMD, CJK, ZYW, MJM, KSO, NVD

Supervision: ELA (Hi-C), SDK (overall)

## Data and materials availability

NCBI: PRJNA613468, PRJNA629833, PRJNA718331, and PRJNA512907. Hi-C: www.dnazoo.org. Code: https://github.com/kocherlab/HalictidCompGen. Genomes and browsers can be accessed at beenomes.princeton.edu. Please address inquiries or material requests to.

## Supplementary Materials

Materials and Methods

Supplementary Results

Figs. S1 to S9

Tables S1 to S15

References (60–148)

