## Supplementary Materials for "Convergent selection on hormone signaling shaped social evolution in bees"

**This PDF file includes:**

Materials and Methods
Supplementary Text
Figs. S1 to S9

**Other Supplementary Materials for this manuscript include the following:**

Table S1. Genome annotation statistics. Species name, total assembly length, karyotype, number of genes included in the assembly, the number of isoforms, and BUSCO scores.
Table S2. Characterization of microRNAs expressed in the brains of each species/population.
Table S3. Correlations between gene age and the proportion of gene sets of interest of each age.
Table S4. TF motif results in promoter regions of social vs. solitary lineages.
Table S5. Orthogroups with significant signatures of selection.
Table S6. GO enrichments among sets of coding sequences with signatures of selection. Parameter provides information on which test was conducted (RELAX or aBSREL), selection type denotes which branches were considered, and/or relevant significance thresholds.
Table S7. Samples included in this study. Sample name, species identification, collection date and location, sex of the individual, library preparation method, BioSample and BioProject accession IDs.
Table S8. Gene expression data for each species and tissue type.
Table S9. Cluster-specific DEGs (FDR < 0.05; 10 PCs, Resolution=0.6) for all 6716 cells in halictid scRNA-Seq experiment
Table S10. Cluster-specific DEGs (FDR < 0.05;6 PCs, Resolution=0.3) for all cells in Cluster 6 (glial cluster)
Table S11. Sequences included in evolutionary analyses for apolpp across insects.
Table S12. Hypergeometric tests examining enrichment of tested gene sets.
Table S13. Wilcoxon rank-sum tests of differences in tissue specificity between the genes of interest and all other genes.
Table S14. Numbers of branches under selection in apolpp across insects.
Table S15. GO enrichments among sets of CNEEs as determined by permutation tests.

#### Methods and Protocols

##### Data generation

###### Sample collections

Details of all samples are provided in Table S7.

###### Genomic DNA extraction

We used Qiagen Genomic Tip (Qiagen, USA, Catalog #10223) kits to extract HMW DNA from bee samples with minor modifications. First, halictids exceed the maximum suggested mass for the kit so individual bees were split into two digestions, one that included both the head and thorax and the other with just the abdomen. For each digestion, bee parts were gently ground with a pestle over dry ice before adding 350 ul of buffer ATL and 4 ul of RNAse A followed by incubation at 37°C for 30 minutes. 50ul of proteinase K and 1 mL of G2 buffer were then added and the samples incubated overnight at 50 C with occasional mixing. The standard genomic tip protocol was subsequently followed, combining multiple digestions from the same sample onto the same column when applicable. Centrifugations were done at 12,000g for 30 and 15 minutes and DNA was eluted in 50 ul of TE.

###### HiC sequencing

For Hi-C scaffolding of the 10X drafts, *in situ* Hi-C libraries were generated as previously described[60]. Specifically, head and thorax tissue from a single bee for each species (with the exception of *H. quadricinctus,* for which a single abdomen was used) was crosslinked and then lysed with nuclei permeabilized but still intact. DNA was then restricted and the overhangs filled in incorporating a biotinylated base. Free ends were then ligated together *in situ*. Crosslinks were reversed, the DNA was sheared to 300-500 bp and then biotinylated ligation junctions were recovered with streptavidin beads. This was followed by a standard Illumina library construction protocol. Briefly, DNA was end-repaired using a combination of T4 DNA polymerase, *E. coli* DNA Pol I large fragment (Klenow polymerase) and T4 polynucleotide kinase. The blunt, phosphorylated ends were treated with Klenow fragment (3' to 5' exo minus) and dATP to yield a protruding 3- 'A' base for ligation of Illumina's adapters which have a single 'T' base overhang at the 3' end. After adapter ligation DNA was PCR amplified with Illumina primers for 8-12 cycles and library fragments of 400-600 bp (insert plus adaptor and PCR primer sequences) were purified using SPRI beads. The purified DNA was captured on an Illumina flow cell for cluster generation. Libraries were sequenced on the Illumina sequencer following the manufacturer's protocols.

###### Whole-genome resequencing

For whole genome resequencing, we used the standard Qiagen DNEASY (Qiagen, USA, catalog #69504) protocol to extract DNA, eluting in 100 ul of buffer AE. Libraries were constructed using the PCR-free NuGEN Ovation Rapid DNA (TECAN, USA, catalog #0328-96) library prep kit and sequenced on the Illumina HiSeq4000 or HiSeq2500 with 100 base paired-end reads.

###### Genome assembly

We built 10x Genomics linked-read libraries for a single individual from each species and, in general, each library was sequenced on approximately half a lane of the Illumina HiSeq X platform with 150 base paired-end reads. *Lasioglossum vierecki* is quite small so to obtain sufficient DNA, we pooled two males together for extraction. For *Augochlorella aurata*, two separate libraries were constructed, each from a different individual. One of these libraries was sequenced on a half-lane and the other was sequenced on a full lane. *Lasioglossum figueresi* was sequenced on two, half-lanes, and *Halictus rubicundus* and *Agapostemon virescens* were each sequenced on a single full lane. While most samples used were diploid females, we used single haploid males for *A. aurata, L. oenotherae, L. pauxillum,* and *L. leucozonium.* A single additional individual from each species was also collected for Hi-C library preparation except for *H. quadricinctus,* which is large enough that the same bee was split for Hi-C (abdomen) and 10x (head).

###### Transcriptome sequencing for genome annotation

For RNA-sequencing, heads, abdomens, and most larvae and pupae were extracted using Zymo Quick-RNA extraction kits (Zymo, USA, cat. #R1057). For antennae, Dufour's glands, and very small larvae, the

ThermoFisher PicoPure RNA Isolation Kit (ThermoFisher, USA, cat. #KIT0204) was used. Libraries were made using the NEBNext Ultra Directional RNA Library Prep Kit (NEB, USA, cat. #E7760) with Poly-A bead enrichment and were sequenced on two lanes of the Illumina HiSeq4000 with 100 base paired-end reads. Antennal libraries from *L. zephyrus* and *L. oenotherae* as well as all Dufour's gland libraries were prepared using the NEBNext Ultra II RNA Library Prep Kit (NEB, USA, cat. #E7770) with Poly-A bead enrichment and were sequenced on one Illumina HiSeq4000 lane with 150 base paired-end reads.

We collected transcriptomic data from 11 species (*Ag. virescens, Augochlora pura, Augochlorella aurata, H. ligatus, L. calceatum, L. figueresi, L. leucozonium, L. malachurum, L. marginatum, L. pauxillum,* and *L. vierecki*). For each species, we constructed separate RNA-seq libraries from whole abdomens, whole heads, and antennae, and each library was sequenced to a depth of approximately 10 million reads. For *L. pauxillum* and *L. calceatum* we also sequenced larvae and pupae. Antennae were additionally sequenced from *L. zephyrus* and *L. oenotherae*, and additional antennal libraries were prepared for *A. pura* and *L. marginatum.* Dufour's glands were extracted and sequenced from a total of 12 species (*A. aurata, A. pura, L. leucozonium, L. marginatum, L. vierecki, L. zephyrus, L. calceatum, L. albipes, L. malachurum, L. oenotherae, L. pauxillum,* and *L. figueresi*). These four antennal and all Dufour's gland libraries were sequenced to a depth of approximately 17 million reads.

Brain single-cell RNA-Sequencing: Brain dissection and cell recovery

Four single-cell RNA-Sequencing (scRNA-Seq) experiments were prepared from whole brains of *Lasioglossum zephyrus* and *Lasiogossum albipes*. For *L. zephyrus*, three queen or worker brains were dissected and pooled per sample, and for *L. albipes* a single worker brain was used for each of two samples. We followed the "short dissociation protocol" from [61]. Briefly, brains were dissected in ice-cold Dulbecco's PBS (DPBS) and transferred to 500 µl DPBS containing 0.05% Trypsin-EDTA. Manual cell dissociation was performed via shaking at 25ºC, 1000 rpm for 20 min with gentle trituration every 5 min. Cells were then washed with cold DPBS and resuspended in 400 µl of DPBS with 0.04% BSA and passed through a 40 um Flowmi® cell strainer. Cell counting was performed using propidium iodide-based sorting on the MoxiGO system (Orflo Technologies, Ketchum, ID USA).

#### miRNA isolation and sequencing

miRNAs were isolated from flash-frozen bee brains using the mirVana miRNA isolation kit (AM1560, ThermoFisher). Libraries were constructed using NEBNext Small RNA Library Prep Kit (E7330S, NEB) and sequenced on an Illumina HiSeq, single-end 50 bp reads.

#### JH extraction and LC-MS

Bee brains and other tissues were flash frozen in liquid nitrogen, then ground and centrifuged before adding 200 µl of extraction solvent (40:40:20 methanol:acetonitrile:H2O with 0.5% formic acid[62]) supplemented with 0.2 µg/ml valine-D8 as an internal standard and 16.8 µl of 15% ammonium bicarbonate. LC-MS was carried out following previously described protocols[63]. LC was performed using a Xbridge BEH amide HILIC column (Waters) with a Vanquish UHPLC system (Thermo Fisher). Solvent A was 95:5 water: acetonitrile with 20 mM ammonium acetate and 20 mM ammonium hydroxide at pH 9.4. Solvent B was acetonitrile. The gradient used for metabolite separation was 0 min, 90% B; 2 min, 90% B; 3 min, 75%; 7 min, 75% B; 8 min, 70%, 9 min, 70% B; 10 min, 50% B; 12 min, 50% B; 13 min, 25% B; 14 min, 25% B; 16 min, 0% B, 21 min, 0% B; 21 min, 90% B; 25 min, 90% B. MS analysis was performed using a Q-Exactive Plus mass spectrometer (Thermo Fisher) in positive ionization mode. An m/z range of 70 to 1000 was scanned. Measured JH in *Bombus impatiens* is presented in Figs. S6-S8. Chemical structure of JH III, parent and fragment m/z values for labeled and unlabeled forms, chromatograms and absolute quantification of JH III and JH III-D3 are shown in Fig. S8.

#### Stable isotope tracing

Bumblebee (*Bombus impatiens*) workers were collected from a queenright colony (Koppert Biological Systems, Natupol Excel) and isolated for 24 hours. *Augochlorella aurata* foundresses were collected in May 2022 from Princeton, NJ and were fed sucrose solution overnight in isolation. Following isolation, bees were chill anesthetized until immobile, and then treated with 50ug (*B. impatiens*) or 25ug (*A. aurata*) of deuterated JH III (JH III-D3) (Toronto Research Chemicals, E589402) dissolved in acetone or acetone alone as a control. Treatments were applied by pipetting 1-2 ul of 25 ug/ul JH III-D3 onto the abdominal sternites. Bees remained on an ice pack until the treatments were absorbed and dry. Then, they were returned to isolated tubes for 2, 4, 8, 16, 24, or 72 hours until frozen. For LC-MS of *B. impatiens*, bees were chill anesthetized and then decapitated. For LC-MS of *A. aurata,* heads were lyophilized at -80C and 300 mTorr for 30 minutes, after which brains were dissected and stored at -80°C until analysis.

**Data Analysis**
Code for analyses presented below is available at https://github.com/kocherlab/HalictidCompGen.

#### Genome assembly

Initial assemblies showed the presence of substantial numbers of Illumina adapter sequences. We therefore performed stringent adapter removal. Adapter trimming was performed by running cutadapt v1.11 [64] twice, both times using the adapter sequences AGATCGGAAGAGCGTCGTGTAGGGAAAGAGTGTAGATCTCGGTGGTCGCCGTATCATT, AATGATACGGCGACCACCGAGATCTACACTCTTTCCCTACACGACGCTCTTCCGATCT, and TGTAGATCTCGGTGGTCGCCGTATCATT. The first time, the defaults were used to only remove three base matches to adapters from the ends of both the forward and reverse reads (-a, -A). The second time, adapters were removed from within reads (-b, -B), requiring matches of at least four bases. Reads with any sign of adapter sequence were removed from the dataset rather than being trimmed.

Assemblies on the resulting adapter-filtered data were then performed using Supernova version 1.0.0 [65]. Note that we attempted to use later versions but, while successful on some species, the program crashed on others so we settled on 1.0.0 for the sake of consistency. Each species was assembled three times using random subsets of 105 million read pairs. Initial explorations of the coverage needed using *L. albipes* suggested that ~105 million reads maximized assembly quality metrics. Genome files were then generated using the mkfasta command with style pseudohap and minimum scaffold size of 1,000 bases. These three assemblies were then sequentially merged using quickmerge[66] in every possible order of all three and also subsets of just two assemblies. This yielded a complete set of 27 assemblies, including the original three. Each assembly was then evaluated based on continuity statistics (e.g. N50 lengths and numbers), as well as completeness as determined by BUSCO2[16,67,68] using *Apis mellifera* as the seed species and the set of 4,415 Hymenoptera genes from OrthoDB v9. Gaps were then closed with the sealer module of ABySS 2.0[69] using the settings -B 3000, -G 1000, -P 10, though for certain genomes, these parameters had to be decreased to keep the runtime under reasonable limits.

*L. figueresi* has a larger genome than other *Lasioglossum* so the 10x library was sequenced on two half lanes. The data were filtered as for other genomes and then all data were used to make the assembly. No random subsets were used and no merging was performed as it did not improve the assembly. Similarly, gapclosing with sealer was not performed for this genome because it did not substantially alter the resulting assembly.

*Augochlorella aurata* also has a much larger genome than most other species examined. Two 10x libraries from one haploid male and one diploid female were constructed. One was sequenced on a half lane of Illumina HiSeqX and the other on a full lane of HiSeqX. All data was pooled for analyses. Assemblies using the approach outlined above were of poor quality. FASTQC v0.11[70] analysis of the raw data revealed that the libraries were highly GC-biased with overrepresentation of tandem duplications of the motif ATGGGCT. We therefore filtered 99% of reads containing two tandem sequences of this motif. We kept 1% of these reads assuming that they were being overamplified in the library but still represented real sequence in the genome. This procedure substantially improved the contiguity of the assembly but we suspected that additional data may be present in the 10x libraries that may have been filtered out before assembly. We therefore also ran fragScaff[71] using a half lane of one of the sequencing libraries. We aligned the data with LongRanger[72] and then performed fragScaff analysis with the parameters -b 1 -E 3000 and -G R which approximately doubled the N90 size of the assembly.

A similar procedure removing overrepresented motifs was also required for *H. rubicundus* (motif TATCCTATCC), *H. quadricinctus* (motif TATCCTATCC), and *Ag. virescens* (motif TAACCCAGACC). Additional scaffolding with fragScaff was performed for *H. rubicundus* but not the other two species as it was not found to improve the assembly.

#### Hi-C scaffolding

We sought to perform Hi-C scaffolding to error-correct, order and orient as well as anchor the draft genomic sequences to chromosomes[15,73]. For this, candidate chromosome-length genome assemblies were generated using 3D-DNA[15] followed by additional finishing using Juicebox Assembly Tools[15]. The resulting contact matrices as well as matrices corresponding to the draft assembly, before the Hi-C scaffolding, are available on www.dnazoo.org.

#### Transcriptome assembly

We generated three directional RNAseq libraries from a single individual per species. Tissues were: antennae, heads, and abdomens from *A. virescens, L. figueresi, A. pura, L. marginatum, A. aurata, L. leucozonium, H. ligatus, L. pauxillum, L. vierecki, L. malachurum,* and *L. calceatum.* For each species, multiple libraries were combined and assembled using Trinity[74] with the –trimmomatic, --SS_lib_type RF and –normalize_reads options. These libraries were also assembled using the genome-guided approach by first mapping to the genome assemblies using HISAT2 v2.0.5[75] and setting a maximum intron length of 100,000 bases.

The non-directional libraries from the antennae of additional individuals from *A. pura, L. marginatum, L. zephyrus,* and *L. oenotherae* and the Dufuor's glands from *A. aurata, A. pura, L. leucozonium, L. marginatum, L. vierecki, L. zephyrus, L. calceatum, L. albipes, L. malachurum, L. oenotherae, L. pauxillum,* and *L. figueresi* were assembled separately from the directional libraries using both *de novo* and genome-guided Trinity and the –trimmomatic and –normalize_reads options.

For *L. albipes*, we assembled two datasets in addition to the Dufuor's gland. First, we took the RNAseq data from an existing *L. albipes* paper[76] and reassembled it using *de novo* and genome-guided Trinity. We also single-end sequenced two directional libraries constructed from the head and antennae of a single individual.

Normalized gene expression data for each species and tissue type is provided in Table S8.

#### Filtering bacterial scaffolds

Initial genome assemblies included varying numbers of scaffolds of likely bacterial origin. We filtered these by comparing scaffolds to previously sequenced genomes. First, the database of complete bacterial genomes from NCBI was downloaded on February 7, 2017. We then used BLASTN to compare all assembled scaffolds to the bacterial genome database using default parameters. Significant hits to bacterial genomes were merged if they were within 10 bases of each other. If more than 25% of the length of a scaffold had a significant hit to a bacterial genome or more than 10% of the length of the scaffold in more than 10 segments had a significant hit to a bacterial genome then the scaffold was extracted. These sequences were then compared with a separately downloaded full set of animal genomes from NCBI using BLASTN, again with default parameters. If a region had a higher percent similarity to bacterial genomes than animal genomes then that scaffold was considered of bacterial origin and was removed from the assembly prior to HiC scaffolding and annotation.

#### Repeat masking

In order to obtain high quality genome annotations, we first characterized the repetitive elements present in each individual genome using RepeatModeler v1.0.7[77]. We removed redundancy from the resulting set of repetitive elements by clustering those with 80% or greater similarity using CD-HIT v4.6[78]. We compared the resulting repeat elements to all UniProt proteins and *Drosophila melanogaster* proteins using BLASTX. Putative repetitive elements with bitscores of at least 100 or more than 50% similarity over 50% of the length of either the protein or repeat sequence were removed from the set of repetitive elements so as to avoid masking protein sequences in the genome. Only repetitive elements longer than 80 bases were retained.

The resulting set of repeat elements was combined with the repetitive elements for Arthropods available from RepBase (downloaded from https://www.girinst.org/ on March 8, 2017) and then used to mask each genome.

#### Identification of conserved non-exonic elements (CNEEs)

Repeat-masked genomes of each of the 19 species included in our study were aligned using Cactus[17,79]. We followed the methods outlined in Sackton et al.[80] (https://github.com/tsackton/ratite-genomics/tree/master/04_wga) to identify conserved non-exonic elements. First, fourfold degenerate sites were extracted from the alignment based on the *Nomia melanderi* annotation using halTools v2.1. phyloFit, part of the PHAST package[81], was used to generate a model of neutral evolution across these species using these data. We generated five of these neutral models using different random starting points and all models were nearly identical so one was chosen arbitrarily. This model was updated to reflect the average GC content in extant genomes using modFreqs.

We used phastCons[82] to identify conserved non-exonic elements. First, MAFs (Mutation Annotation Format files) were generated from the Cactus HAL files using *Nomia melanderi* as a reference sequence.

Self-alignments from other regions of the *N. melanderi* assembly were filtered out of each aligned block and the MAFs were filtered using mafDuplicateFilter from mafTools[83]. mafStrander was then used to force all alignment blocks to a positive strand for the *N. melanderi* reference. Finally, alignment blocks were sorted based on coordinates using mafSorter and split into non-overlapping 1Mb blocks with at least 1,000 informative sites.

We ran phastCons using a range of parameters to identify optimal settings for identifying CNEEs. In total, we ran the procedure nine times with every combination of expected conserved element length of 20, 45, and 70 combined with target coverage proportions of 0.2, 0.3, and 0.4. Based on these results we chose a conserved element length of 45 and target coverage of 0.3 for identifying conserved elements. *N. melanderi* coding sequences were removed from the resulting conserved elements. We required that a conserved element be at least 100 bases in length in *N. melanderi* to be included in our dataset. The coordinates of the resulting CNEEs were identified in each species using halLiftover. Coding sequences were then removed from the CNEEs in each species individually and sequences of less than 100 bases were discarded. CNEEs were aligned with FSA[84] and alignments were filtered using the "-automated1" option of TrimAl v1.4.rev22[85].

#### Analysis of CNEEs

In order to identify CNEEs associated with social evolution, we examined differences in the rates of evolution in CNEEs among extant taxa. For each locus, branch lengths were calculated on the species topology using BASEML v4.9[86,87]. The resulting branch lengths were standardized to average rates of evolution in each genome using the branch lengths estimated by RAxML[88] using fourfold degenerate sites extracted using CODEML v4.9[86,87]. We then compared the standardized branch lengths of the five pairs of closely related social and solitary species in our study (*Augochlorella aurata* vs. *Augochlora pura*, *Halictus ligatus* vs. *Halictus quadricinctus*, *Lasioglossum marginatum* vs. *L. figueresi*, *L. zephyrus* vs. *L. vierecki*, and *L. pauxillum* vs. *L. oenotherae*). Loci for which the branch lengths were longer in all social taxa than the most closely related solitary taxa were considered fast-evolving in social species and vice versa.

#### miRNA annotation

miRNA was extracted from whole brains of individual females using the *mir*Vana™ miRNA Isolation Kit (Invitrogen cat. no. AM1560), and libraries were prepared with the NEBNext Small RNA Library Prep Set for Illumina (NEB cat. no. E7330S) (sample details in Table S7). We used miRDeep2 (v. 2.0.0.8)[89] to identify miRNAs expressed in the brain of each species. We removed adapters, removed reads shorter than 18 nucleotides or with noncanonical letters prior to mapping reads to the genome assembly using the mapper.pl function. We then used miRDeep2.pl to detect known and novel miRNAs in each species. Known miRNAs were mature miRNA sequences previously described in other species (*Apis mellifera*, *Drosophila melanogaster*, *Nasonia vitripennis*, *Tribolium castenum, Bombyx mori, Bombus impatiens, Bombus terrestris, Megalopta genalis, Megachile rotundata, Nomia melanderi*)[90–92]. We quantified expression with the quantifier.pl script. We then filtered novel miRNAs based on the following criteria: no rRNA/tRNA similarities, minimum of five reads each on mature and star strands of the hairpin sequence, and a randfold p-value < 0.05; randfold describes the RNA secondary structure of potential precursor miR (pre-miRs). To identify homologous miRNAs across species, we added this filtered set to the list of known miRNAs and repeated the miRDeep2.pl detection step, followed by filtering. We characterized the genome location of each miRNA in the genome using bedtools (v. 2.27.0)[93] intersect to find overlap between miRNAs and predicted gene models.

We used computational methods to predict the gene targets of each miRNA in each species. First, we extracted potential target sites 500 bp downstream from each gene model using bedtools flank and getfasta. We used miRanda (v. 3.3)[94] with a minimum energy threshold of -20 (-en -20), minimum score of 140 (-sc 140), and strict alignment to seed region (-strict). We also used RNAhybrid (v. 2.12)[95] with a minimum free energy threshold -20 (-e -20). We kept only those miRNA-target pairs that were predicted by both programs at p < 0.001.

#### Coding sequence annotation

We generated gene predictions for each genome using BRAKER v2.1.0[96,97]. First, all RNAseq reads for each genome were mapped to the repeat-masked genomes using HISAT v.2.0.5[75] with a maximum intron length of 100,000. BRAKER was run on these mapped reads and the repeat-masked genome (softmasked). For *Halictus quadricinctus* (HQUA), for which we were unable to obtain RNAseq data, we used the RNAseq data from *Halictus rubicundus* (HRUB) for BRAKER prediction.

MAKER v3.00.0[98,99] was run on the repeat-masked genomes. The GFF files of aligned ESTs from PASA[100,101] were used as EST evidence. All high-quality protein predictions from Transdecoder[102] from all species were combined and used as protein evidence for each genome. In addition, OGSv3.2 from *Apis mellifera*, OGSv5.42 from *Lasioglossum albipes*, and all UniProt proteins downloaded on December 2, 2016 were included as protein evidence.

All gene predictions from BRAKER were fed to MAKER as pred_gff and we also used the MAKER implementation of EvidenceModeler. We used both the always_complete and correct_est_fusion options in MAKER.

Included in the output from MAKER is a set of gene predictions for which protein and transcript evidence did not overlap (*.non_overlapping_ab_initio.*). We used InterProScan v5.21-60.0[103] to identify those sequences within this set with conserved protein domains. We considered anything that InterProScan assigned an InterPro family to as a true gene and reran MAKER, incorporating these predictions into the annotation. The resulting annotations were used as the basis for the OGS. The only subsequent processing that occurred was the removal of any annotations on scaffolds deemed to be spurious duplications in the assembly step.

An Official Gene Set v2.1 was created for each genome. These gene sets are relatively complete as measured by BUSCO2 when comparing with the 4,415 genes expected to be present in all Hymenoptera species based on OrthoDB v9[104] (Fig. 1). The average percent of complete BUSCOs present in these Official Gene Sets is 93.6%. On average, 2.8% of genes are missing and 3.5% are fragmented.

#### Orthology

We used OrthoFinder v2.3.2[105,106] to identify orthologous groups of genes within the 19 species of Halictidae. We also uploaded protein sets to OrthoDB v10[107]. Gene names, orthologous *D. melanogaster* genes, and orthologous *A. mellifera* genes were assigned based on the OrthoDB groups. If more than 50% of the genes in a particular orthogroup were mapped to the same OrthoDB group, that orthogroup was assigned the gene name from the OrthoDB group.

Gene Ontology terms were assigned to orthogroups using information from three sources. First, Trinotate v3.0.1[108] was used to assign GO terms to every gene in each of the halictid genomes. In addition, GO terms of both the *D. melanogaster* and *A. mellifera* orthologs determined by OrthoDB mapping were compiled for each species. Orthogroups for which at least 30% of the genes in the group had been assigned a particular GO term from at least one of these sources were assigned those GO terms. For GO enrichment results, the find_enrichment.py script from GOATOOLS (v. 1.0.3)[109] was used with our custom orthogroup to GO mapping table. Only Biological Process terms were considered, and Benjamini Hochberg multiple testing correction was used to calculate q-values for GO terms with at least two representative genes in the target set tested for enrichment.

We used the 318 genes identified as transcription factors (TFs) in *Apis mellifera* in the Regulator database[110] to identify TFs in halictids. The *A. mellifera* orthologs determined by OrthoDB and assigned to orthogroups as above were used as the TF set in our study. Using this approach, we were able to identify 295 TFs in halictids.

In order to identify human orthologs associated with autism spectrum disorder (ASD), we identified reciprocal best BLAST hits between the set of human UniProt sequences and the protein sequences from each halictid genome. If at least eight genes in an orthogroup had a reciprocal best hit to humans and greater than half of the genes in an orthogroup were reciprocal best hits to the same human gene, then the orthogroup was assigned to that human gene. Gene associations with ASD were taken from the SFARI Gene web portal (https://gene.sfari.org/) on Oct. 15, 2017. This yielded human gene assignments to 5,787 orthogroups. The SFARI genes are ranked according to the amount of evidence associating them with autism. S is "syndromic" and then the scale is 0 to 6 with 0 having the highest degree of evidence. We included tests of all groups of genes of a certain degree of evidence or greater. We found Halictid orthologs of 30 SFARI "S" genes, 68 of "0", 13 of "1", 29 of "2", 55 of "3", 84 of "4", 41 of "5", and 5 of "6".

#### Gene ages

We used the Phylostratigraphy pipeline (https://github.com/AlexGa/Phylostratigraphy) developed in previous studies[111,112] to identify the approximate evolutionary age of origin for orthogroups in our study. The proteins used as a reference set for identifying gene ages included *Acyrthosiphon pisum* NCBI annotation v2.1, *Athalia rosae* i5k OGS v1.0, *Blattella germanica* i5k OGS v1.0, *Bombyx mori* NCBI Annotation Release 102, *Danio rerio* NCBI Annotation Release 106, *Drosophila melanogaster* FlyBase annotation r6.16, *Helobdella robusta* JGI Helro1 annotation release 46, *Homo sapiens* UniProt Proteome

UP000005640, *Nasonia vitripennis* NasoniaBase OGS v1.2, *Octopus bimaculoides* Metazome v3.0 release, and *Polistes dominula* NCBI Annotation Release 100.

The createPSmap.pl script from Phylostratigraphy (https://github.com/AlexGa/Phylostratigraphy) was run on each of the 19 species included in this study with a sequence offset of 50 and an e-value of $1x10^{-5}$. To assign an age to an orthogroup, we required that representative genes from at least five species be assigned an age by Phylostratigraphy and that the majority of the genes with an assigned age be assigned to the same age.

To test for correlations with ages, we extracted the crown age of taxonomic levels from the literature[113–115]. The ages used were 684 Mya for Bilateria (Age=1), 632 Mya for Protostomia, 373 Mya for Neoptera, 345 Mya for Holometabola, 240 Mya for Hymenoptera, 192 Mya for Apocrita, 162 Mya for Aculeata, 134 Mya for Apoidea, and 71 Mya for Halictidae (Age=9).

#### Gene expression

For each RNAseq sample, we calculated TPM (transcript per million) values for each gene using Salmon v0.9.1[116] with quasi-mapping and controlling for GC bias. Dufour's gland libraries were of the IU form; all other libraries were directional for which we instead used ISR. For all subsequent analyses, we used quantile normalization (normalize.quantiles from the preprocessCore library in R[117]) to normalize TPMs across species for each tissue. We then performed PGLS (phylogenetic generalized least squares) analyses using the R package geiger v2.0[118] to identify correlations between sociality and expression level for each orthogroup, excluding those species with polymorphic social behaviors.

#### Brain single-cell RNA-Sequencing

Raw sequencing reads from two *L. zephyrus* and two *L. albipes* brain scRNA-Seq experiments were trimmed with fastp[119] using default settings and aligned to respective genomes with CellRanger V6 (10X Genomics). Prior to analysis, we converted *L. albipes* gene IDs to one-to-one orthologs in *L. zephyrus* such that each species could be analyzed in a common transcriptomic landscape. We performed normalization and integration to combine scRNA-Seq data across species using the SCTransform pipeline with default settings in Seurat V4[120,121]. This yielded a final dataset of 6,716 cells for downstream analysis. Genes without one-to-one orthologs were retained in the Seurat data object and used for differential expression analysis but not for integration or clustering. This allowed us to identify cell types based on genes shared between the two species while also permitting genes that were both species- and cluster-specific to be identified.

The top 3,000 most variable genes were used to approximate the true dimensionality of the dataset, and a ranking of principle components (PCs) by percentage variance explained indicated that ten dimensions were suitable for cell clustering and visualization. Dimensionality reduction was performed with the Uniform Manifold Approximation and Projection (UMAP) technique, and cell clusters were identified with shared nearest-neighbor Louvain modularity optimization using a resolution of 0.6. Finally, cell-type-specific genes were identified using the MAST algorithm[122] in the "FindAllMarkers" function with "only.pos=T" and a Bonferroni-corrected *P* value of 0.05 (Supplementary Table S9).

Our analysis identified 11 cell clusters that could be broadly categorized as "neurons" or "glia" based on the mutually exclusive expression of either canonical neuronal markers *fne* and *Syt1* or glial markers *repo* and *bdl*, in accordance with previous sc analyses of insect brains[61,123,124]. Upon closer examination, we found that *apolpp* was specifically upregulated in the only glia cell cluster, Cluster 6, along with two lipoprotein receptors, *LpR1* and *mgl*, as well as *apoltp*, all of which have been shown to play a role in neuron-glia lipoprotein shuttling and sequestration[125,126]. The juvenile hormone (JH) epoxide hydrolase (*jheh2*) was also significantly upregulated in Cluster 6 relative to other cell clusters.

To gain deeper insight into the specific expression patterns of *apolpp* and related genes in glia, we subsetted and clustered glial cells using six PCs and a resolution of 0.3 to compensate for the smaller cell population, yielding five subclusters within the glial population. We then identified glial cell-type-specific DEGs following the method described for the total cell population, and this finer-grained analysis revealed that *apoltp* and *apolpp* were co-expressed in a glial subcluster that also expressed *moody*, a marker of subperineurial glia, a subtype of surface glia that form the blood brain barrier (BBB) and regulate its permeability in the *D. melanogaster* brain[46,47]. In a separate subcluster, we found that *jheh2* was significantly upregulated along with *zyd*, a marker of cortex glia[124,127,128] (Supplementary Table S10). Unlike surface glia, cortex glia encase and support neuronal somata and are thought to play a role in juvenile hormone sensitivity[129,130].

#### Tissue specificity

We calculated the specificity of expression of each orthogroup across the four tissues examined using the normalized expression levels[131]. We required data from at least eight species and all four tissues in order to calculate a specificity index for each gene. When multiple genes were included in an individual orthogroup, a single representative was randomly chosen for the specificity analysis. The specificity metric ranges between 0 and 1. The closer the value is to 1, the more tissue-specific the gene.

#### Coding sequence evolution

*Alignment.* Coding sequences were aligned using the codon-aware version of Fast Statistical Aligner v1.15.9[84]. For several analyses, we then performed an extensive filtering procedure to exclude poorly aligned regions. This filtering followed Sackton *et al.* [80] (https://github.com/tsackton/ratite-genomics/tree/master/03_homology/03_protein_coding_alignment). Each site in an alignment was only retained if at least eight sequences and 30% of sequences were a non-gap base at that site and were represented by at least four species. We then masked poorly aligned regions using the scripts of Jarvis *et al*. [132]. Following this masking, the first filter of alignment columns was performed again on the masked data. We required that 50% of the length of the original sequences not be masked by any of these steps and that less than 50% of the aligned sequence be gap characters. All sequences had to have at least 300 non-gap or masked bases.

*Relaxation using HyPhy RELAX.* We used HyPhy RELAX v2.3.11[33] to identify genes experiencing relaxed selection in solitary species that evolved from ancestrally social lineages. For an orthogroup to be tested, we required that at least 12 taxa be present and that *Halictus ligatus, H. quadricinctus, Augochlorella aurata, Augochlora pura,* and at least one pair of *Lasioglossim marginatum* / *L. figueresi, L. zephyrus* / *L. vierecki*, and *L. pauxillum* / *L. oenotherae* be represented. This requirement ensured that at least one closely related pair of social and solitary species from each of the three social clades was included. Socially polymorphic species were excluded for this analysis. 6,904 loci met these requirements. In addition to examining signatures of relaxation in solitary species, we performed a parallel test on the extant lineages of social species to represent a null baseline. We do not expect an elevation of relaxed selection in social species that evolved from social ancestors. FDR-correction was used to account for multiple testing at FDR<0.1.

*Selection using aBS-REL.* We also used HyPhy aBSREL[18] tests to identify signatures of selection on individual branches. To include an orthogroup in this test, we required that at least two *Halictus* species, at least two of *Augochlorella aurata, Augochlora pura,* and *Megalopta genalis*, at least two *Lasioglossum*, and at least two of *Dufourea novaeangliae, Nomia melanderi,* and *Agapostemon virescens* be represented. A total of at least 12 species had to be represented. This test yielded p-values for every branch across every locus. This full set of p-values was FDR-corrected for multiple testing at FDR<0.05.

Because we did not require that all taxa be represented to include a locus in this test, identifying the same branches across loci was not straightforward. Therefore, we stringently defined branches based on the extant taxa that make up the descendants of those branches. We required that a locus include sequences for all descendants of a branch as represented in the full dataset as well as at least one member of the sister clade. In addition to identifying specific branches under selection, we also counted the number of branches identified as under selection in a locus regardless of taxonomic composition.

*Phylogeny inference.* Fourfold-degenerate sites were extracted from the “4fold.nuc” files produced by PAML during inference of ancestral sequences. For those orthogroups with single representative sequences in each of the 19 halictid species, these sequences were concatenated, producing a 561,041 base matrix. The phylogeny of the group was inferred using RAxML with a GTRGAMMA model of evolution. The resulting topology matched previous results[3]. This phylogeny (with the inferred branch lengths) was used in all of the evolutionary analyses outlined above.

#### Associations between promoter TFBSs and sociality

Previous studies examining associations of genetic elements with social evolution in bees found that transcription factor motif binding strength in promoters was more often positively correlated with social behavior in 223 motifs identified in *Drosophila*[7]. We replicated those analyses here with a few minor differences.

For each of the motifs used in that previous study[4], a “stubb” score[22] was calculated for each 500 base window in every genome, with a step size of 250 bases. For each motif for every species represented in each orthologous group, we then identified the window with the greatest stubb score within 5kb upstream and 2kb downstream of the transcription start site and assigned that score to that gene for that species. Then, for a given species and a given motif, the motif scores were rank-normalized (best score=1, worst score=0). We did not perform the GC-normalization and p-value calculation used previously[7], instead analyzing these rank scores directly.

For every orthogroup and every motif, we examined correlations between stubb score and social behavior using PGLS as implemented in geiger v2.0[118] after first removing all socially polymorphic taxa. These analyses were performed on the phylogeny generated by RAxML on a concatenated matrix of all fourfold degenerate nucleotide sites present across all 19 halictid species. In order to consider the correlation of a particular motif and orthogroup with social behavior, we required that the normalized rank of that motif be at least 0.95 in at least one species. Ranks of 0.95 or greater indicate that the stubb score for that particular motif is in the highest 5% among the 7,102 genes considered across the genome, indicating that these sequences are the most likely to actually be recognized and used by TF's to modulate expression. We also required that the number of genes with significant correlations of a motif be at least five. For each motif, we then counted the numbers of significant (PGLS p-value < 0.01) correlations detected with higher stubb scores in social taxa and with higher stubb scores in solitary taxa. Those motifs for which there were differences in numbers of significant correlations between social and solitary taxa of at least 20% were considered to be associated with the behavior with which they more often correlated.

#### Apolpp evolution

*Apolpp within Halictidae.* We explored the evolution of the gene *apolpp* in detail. If the changes in this gene are associated with social behavior, we expect that these changes would change the function of the protein. We therefore attempted to find particular regions of the apolpp protein sequence associated with social evolution. First, we simply examined 100 AA sliding windows with a 50 AA step size across the *apolpp* alignment. For each window, we inferred branch lengths on the species tree using RaxML v7.3.0[88] with the PROTGAMMAWAG model of evolution. We then calculated the branch length between each tip species and the outgroup species, *D. novaeangliae.* Most of the species in our analysis, regardless of social behavior, evolved from a social ancestor. However, three species, *D. novaeangliae, N. melanderi* and *Ag. virescens,* evolved from strictly solitary ancestors. Transitions in social behavior, rather than sociality itself, might be associated with shifts in evolutionary rates, thus we compared the 16 species with any shift in social behavior to *N. melanderi* and *Ag. virescens*. Specifically, we looked for regions with longer branches in all 16 species with at least one ancestral shift in social behavior relative to the two species without such shifts. We also used fpocket2[133], as implemented in the Phyre2[134,135] Investigator tool, to predict the likely active sites of *apolpp* in halictids, using the *H. ligatus* protein sequence as a reference.

*Apolpp across insects.* We also explored *apolpp* evolution across insects as a whole in order to place the patterns of evolution within Halictidae into a broader context. We collected *apolpp* sequences from 78 additional Neopteran insect species. In particular, we included an additional 12 species of bees, 18 species of ants, eight other Hymenoptera, 10 Coleoptera, 12 Diptera, five Hemipterans, and 11 Lepidoptera (Table S11).

Protein sequences were aligned using MUSCLE v3.8.31[136,137] and filtered (with the -automated1 option) and backtranslated to nucleotides with trimAl v1.4.rev22[85]. We ran aBSREL as implemented in HyPhy v2.3.11 on the resulting nucleotide alignment, testing all 191 branches within the phylogeny. We used FDR-correction to adjust the resulting p-values and counted the number of branches detected as experiencing selection in each of the major clades examined.

We also examined shifts in rates of protein evolution during the evolution of Halictidae. We performed likelihood ratio tests assessing the possibility of rate shifts between five pairs of sister taxa: Holometabola versus Hemiptera, Hymenoptera versus all other Holometabola, ants and bees versus other Hymenotpera, bees versus ants, and Halictidae versus other bees. For each test, we used PAML to calculate the likelihoods of models of evolution that fixed the rates of evolution in the two clades being compared, or allowed them to differ. We also repeated the same set of LRTs as above using just the 51 amino acids present in the pockets predicted by fpocket2.

Finally, we used the branch model of PAML v4.9[86,87] to estimate dN/dS ratios for each of the major clades examined.

#### LC-MS Data Analysis

Data were analyzed using El-MAVEN Software (Elucidata, LLC., elucidata.io). Chromatographs and mass spectra were exported from El-MAVEN and plotted using MATLAB, and peak areas were exported, processed and plotted using R. Peak areas were all normalized to a valine-D8 (Cambridge Isotopes Laboratories Cambridge, MA) internal standard. Absolute quantification of JH III and JH III-d3 was carried out using a standard curve of purified JH III (Sigma, Cat. #J2000) and JH III-d3 (Toronto Research Chemicals, Cat. # E589402), respectively. For social *A. aurata* and solitary *A. pura* comparisons, JH III quantities were standardized to ng/mm3 using brain volumes previously-quantified for each species[138]. Raw data available upon request.

### Supplementary Text

#### Coding sequence evolution

*Signatures of selection across halictids.* We tested 7,769 genes for signatures of positive selection within Halictidae using HyPhy aBS-REL. We obtained p-values for every branch present in the phylogeny for every gene. Thus, we only examined those p-values that passed FDR correction, combining all p-values across all loci and branches for correction. Most basically, we examined those genes with signatures of selection on at least a given number of branches ranging from one to 10. For this analysis we were agnostic to the taxonomic composition of those nodes identified as under selection (i.e., clades did not have to be fully represented taxonomically to be included in these gene sets). Finally, we also examined all genes identified with signatures of selection on all 35 branches present in the complete phylogeny of all 19 species.

4,261 genes show signatures of selection on at least one branch in the phylogeny (FDR-corrected $p < 0.05$; Table S5) which are enriched for three GO terms (Fisher's Exact $q < 0.1$; Table S6). These GO terms are associated with structural functions (i.e., "cell adhesion", "biological adhesion", and "cell junction assembly"). Genes with larger numbers of branches identified as experiencing differential selection are enriched for larger numbers of GO terms, with the greatest number of GO terms (230) identified in the set of 40 genes with at least nine branches identified as under selection. These terms include "synaptic transmission, glutamatergic" as well as a number of others related to gene regulation and synapse function (Table S6). Previous studies have identified a conserved set of genes associated with social behavior in insects and vertebrates, including ASD in humans[2,139]. We thus looked for overlap between our candidate gene lists and the SFARI gene list, which is a curated list of genes associated with ASD. Many of the genes experiencing positive selection in halictids are also significantly enriched for SFARI genes (Hypergeometric test $p < 0.05$; Table S12). While these significant overlaps do not reveal anything about the differences between social and solitary species specifically, they do indicate that these genes have experienced changing selection pressures within Halictidae as a whole, a highly socially variable family. Interestingly, the gene *Syx1A*, which has been previously implicated in social evolution in Halictidae[2], is identified by aBSREL on six branches within the family (*N. melanderi, L. vierecki, L. marginatum, L. albipes*, the branch ancestral to *L. albipes* and *L. calceatum,* and the branch ancestral to *L. oenotherae* and *L. pauxillum*), supporting the possibility that changes in this gene are fundamental to social behavior within this group.

There is a weak but significant correlation between the number of branches identified with signatures of selection for a gene and the age of that gene (Pearson correlation $p=4.3\times10^{-5}$, $r=0.047$), indicating that younger genes tend to be more malleable. There are also significant positive correlations between the proportion of genes of a given age with at least three ($p=0.023$, $r=0.78$) and four ($p=0.044$, $r=0.72$) branches under selection and the age of those genes, again indicating that younger genes are more likely to be undergoing selective changes (Table S3). Concordantly, genes with any number of branches identified by aBSREL have significantly higher tissue-specificity than other genes (Wilcoxon $p < 0.005$; Table S13).

*Positive selection associated with the origins of eusociality.* We selected the two branches on which social behavior convergently originated within halictids[3]: at the root of the Augochlorini and at the root of the genera *Halictus* and *Lasioglossum*. The aBSREL tests identified 309/6312 genes under positive selection on the Augochlorini origin branch and 62/5058 genes under positive selection on the *Halictus* + *Lasioglossum* origin branch. Of the 4,341 genes that were tested by aBSREL on both of these branches, 191 were significant on the Augochlorini branch and 53 were significant on the *Halictus* + *Lasioglossum* branch; nine genes were detected as experiencing positive selection on both, significantly more than expected by chance (Fisher's Exact, p-value=0.004, 3.97-fold enrichment). Perhaps due to the small number of genes identified, no discernable GO enrichment was found for this gene set, although we do detect significant enrichment for genes that differ significantly in expression in the antennae of social and solitary bees (Table S12).

*Positive selection associated with the loss of eusociality.* We did not find any genes that showed a signature of positive selection on all of the branches associated with the loss of eusociality, but there were 95 genes with evidence of positive selection on at least 1 secondarily solitary lineage in both the Augochlorini and the Halictini *s.s*. These genes are not enriched for any GO terms, but include the JH co-receptor, *taiman* (Table S5).

Intensification of selection on extant eusocial branches.

To search for signatures of selection associated with the maintenance or elaboration of eusociality, we identified genes experiencing intensified purifying or positive selection on terminal eusocial branches in our phylogeny. We found 393 genes with intensified selection on terminal eusocial branches (HyPhy RELAX, FDR<0.1), 15 of which also experienced positive selection on at least one eusocial origin branch (Table S5). These 393 genes are enriched for cell adhesion (GO:0007155, Fisher's Exact q=3.86e-3, enrichment=2.49) and the non-canonical Wnt signaling pathway (GO:0035567, Fisher's Exact q=8.87e-3, enrichment=8.09), among others (Table S6). In addition, genes under intensified selection on eusocial branches included 16 TFs, including *ultraspiracle* (*usp*) and *Ecdysone Receptor* (*EcR*), both of which are associated with the *usp* binding motif previously linked to eusocial evolution in stingless bees[7] and are known transducers in the JH signaling pathway.

Relaxed selection in lineages that have lost social behavior.

Our species sampling includes six species that are solitary but that evolved from a social ancestor (*Augochlora pura, H. quadricinctus, L. leucozonium, L. figueresi, L. oenotherae,* and *L. vierecki*). We assumed that the evolutionary loss of social behavior would lead to relaxed selection in those genes most constrained among social species and for which the fewest changes can be tolerated. We used HyPhy RELAX to identify those genes experiencing relaxed selection across all six of these species, testing 6,890 loci and identifying significant signatures of relaxation in 443 loci (FDR-corrected LRT p < 0.1). Four GO terms were significantly enriched in this gene set, including "chromosome condensation", "DNA packaging", and "DNA conformation change" (Table S6), all three of which may be related to the regulatory functions identified in the correlations between social behavior and constraint.

In order to determine whether the loss of social behavior did, indeed, lead to additional relaxed selection, we also used HyPhy RELAX to identify those genes experiencing relaxed selection in the six extant social lineages represented in our dataset, testing a total of 6,887 loci, 305 of which showed significant signatures of relaxation (FDR-corrected LRT p < 0.1). These 305 genes were enriched for antibiotic metabolic process (GO:0016999, Fisher's Exact, q=0.053, enrichment=4.36), drug catabolic/metabolic processes (GO:0042737, GO:0017144, Fisher's Exact, q=0.093, enrichment=4.03,2.25), and malate metabolic process (GO:0006108, Fisher's Exact, q=0.095, enrichment=11.29) (Table S6). This is a significantly lower proportion than the 443 of 6,890 loci identified to be experiencing relaxed selection among the solitary species (Fisher's exact test $p=2.42 \times 10^{-7}$, odds-ratio=1.48), suggesting that the loss of eusociality is more often associated with a release of constraint compared with eusocial maintenance or elaboration.

The proportion of genes undergoing relaxed selection in solitary species is correlated with gene age; a larger fraction of younger genes is experiencing relaxed selection (Pearson correlation p=0.002, r=0.869; Table S3; Fig. 2). Genes experiencing relaxed selection in solitary species also have significantly greater tissue specificity than other genes (Wilcoxon test p=0.00167; Table S13). Therefore, those genes associated with social evolution tend to be more recently evolved with more tissue-specific function making them less likely to be essential for overall organism function when expressed in a solitary phenotype.

#### Gene ages and patterns of selection

We examined the relationship between gene age and selection associated with eusocial origins, maintenance, and reversions to solitary life histories to determine whether or not there was any relationship between selection and gene age (Fig. S4). There is a greater proportion of young genes experiencing relaxed selection when eusociality is lost (Pearson's r=0.869, p=0.002); we found no significant association with relaxation on extant, eusocial branches. Next, we looked at the sets of genes that show intensification of selection pressures (HyPhy RELAX, FDR<0.1), but neither of these sets showed any significant association with gene age. Finally, we looked at genes that showed evidence of positive selection (HyPhy aBSREL, FDR<0.05). We found no relationship between gene age and the proportion of genes with evidence of positive selection on at least 1 branch representing the origins of eusociality (Positive, gains). Likewise, there was no relationship between gene age and the proportion of genes with evidence for positive selection (HyPhy aBSREL, FDR<0.05) on at least 1 loss branch in the Halictini and on the 1 loss branch in the Augochlorini (Positive, losses). Interestingly, when we combine the lists of genes with signatures of positive selection on the Augochlorini (309) and Halictini (62) branches, we find a trend suggesting a higher proportion of younger genes experience positive selection (linear regression, $R^2$=0.415, p=0.0846). These results add some support to our observation that younger genes are more likely to experience relaxed selection when eusociality is lost.

#### Complementary patterns of selection associated with emergence and breakdown of eusociality

There were 4,334 loci for which we tested the two social origin branches for signatures of positive selection and that we tested for signatures of relaxation on extant solitary species with social ancestors using HyPhy RELAX. Of the 443 genes detected as experiencing relaxed selection, four of them overlap both sets of genes with signatures of positive selection on both origin branches. This is a significant enrichment of overlapping signals between these three tests (Multi-Set Enrichment, enrichment=8.85, p-value=0.001). Given the overlap of these signals, these genes (*apolpp*, *Pat1, Hex110*, and a Hymenoptera-specific gene, OG_11519) are likely involved in the evolution of sociality. We were unable to assign any gene ontology terms to OG_11519, indicating that it is either completely novel or has diverged substantially in sequence from previously characterized protein domains.

#### Positive selection associated with origins and intensification of selection on extant eusocial lineages

We find 15 genes that show a signature of positive selection on at least one of the branches associated with the origins of eusociality and intensification of selection on extant eusocial branches (14 Augochlorini, 1 Halictini, 0 shared), suggesting that these 15 genes may be particularly relevant to the emergence and elaboration of eusociality (Table S5). 14/15 of these genes are conserved across Bilateria, suggesting that this set is highly conserved and consistent with regression results on the 393 genes experiencing intensification of selection above. Some of the more interesting genes in this set include: *Nop60B* (RNA metabolism), *Bruce* (an inhibitor of apoptosis), and *ReepA* (ER stress prevention in the presynapse). These genes are also enriched for embryo implantation (GO:0007566, Fisher's Exact, q=9.92e-3, enrichment=153; Table S6).

#### Intensification in social lineages, Relaxation in solitary lineages

34 genes show intensification of selection on extant eusocial lineages and relaxation in secondarily solitary species (HyPhy RELAX, FDR<0.1 for both tests; Table S5). These genes likely represent trade-offs associated with the maintenance or elaboration of eusociality, and they are enriched for regulation of SNARE complex assembly (GO:0035542, Fisher's Exact, q=0.074, enrichment=80.91; Table S6), which is a key component of synaptic transmission that has also been implicated with variation in social behavior in *L. albipes*[2] and wasps[35]. None of the 34 genes showed a signature of positive selection on either of the branches representing the origins of eusociality, nor do they show a correlation with gene age.

#### Overlap with previous comparative genomic studies in bees

We did not find a significant overlap between our gene lists and those presented in [7]. We used *L. albipes* reciprocal blasts to translate from the list of genes identified as under selection for Halictidae origins[7] to our new *L. albipes* annotation, which results in some loss of genes. Of the 105 genes remaining after translation (167 in [7]), none overlapped with the genes we identified as under selection in Halictini (not different from random expectation; p=0.487), and 5 overlapped with genes we identified under selection in Augochlorini (not different from random expectation; p=0.224). Five genes (1 of which is also under selection in Augochlorini) that are experiencing relaxed selection on solitary branches overlapped with the genes identified in [7] , but this is also not significantly more than expected by chance (p=0.494). This suggests that our study design and approach is identifying novel mechanisms associated with the evolution of eusociality relative to those previously identified using more distantly related species[7].

#### Apolpp evolution within halictids

We identified seven 100-residue regions evolving faster in lineages with at least one ancestral transition in social behavior than in *Ag. virescens* and *N. melanderi.* Five of these regions occurred within the first 900 amino acids of *apolpp,* comprising a total of 400 amino acids. The vitellogenin-N domain of *apolpp* makes up residues 41-573 and is likely the receptor-interacting site[140]. This region encompasses two overlapping windows that have evolved faster in lineages with social transitions from residues 200-300 and 250-350, indicating that these windows are, indeed, likely to be functionally significant. 51 sites were predicted to be part of the protein pockets of this gene, all of which are within the first 950 residues and 17 of which are also within those windows found to be evolving faster on lineages with social transitions. We also performed the same rate shift analysis using just the residues identified as a part of the pocket of *apolpp* and, indeed, rates of change are higher in lineages with social transitions.

#### Apolpp evolution across insects

We tested five nodes in the phylogeny of *apolpp* across insects for rate shifts and four of these branches showed significant differences in rates (LRT p < 0.01; Fig. S9). Rates appear to have increased significantly in Holometabola as a whole, to have decreased in Hymenoptera relative to the rest of Holometabola, and then increased in ants and bees relative to other Hymenoptera, and again in Halictidae relative to other bees. Rates did not shift significantly between bees and ants (p=0.979). Locations of rate changes are indicated in Fig. S9. Tests of the same clades but with an alignment limited to only those 51 amino acids predicted to be a part of the protein pocket are all nonsignificant (LRT p > 0.05) except for the test between ants and bees and the rest of Hymenoptera, which shows a significant rate increase within the ants and bees (p=0.0051).

We also used HyPhy aBSREL to identify those branches with signatures of positive selection for *apolpp*. Remarkably, 32% of branches within Halictidae have experienced positive selection, a larger proportion than in any other clade examined (Table S14). dN/dS ratios were also highest for halictids (0.19) compared with all other clades examined (Coleoptera (0.021), Hemiptera (0.062), Lepidoptera (0.088), Diptera (0.096), ants (0.17), and non-halictid bees (0.14)). *Apolpp* appears to change particularly quickly within Halictidae. Results are summarized in Fig. S9.

#### TFBS evolution in promoters

To assess the degree to which changes in gene regulation may facilitate the evolution of social behavior in halictids, we characterized TF motifs in putative promoter regions in each halictid genome. For each species, we defined these regions as 5kb upstream and 2kb downstream of the transcription start site for each gene[7] and calculated a score for a given TF in each region that reflects the number and/or strength of predicted binding sites[22]. For each of the 223 motifs examined, we counted the numbers of significant (PGLS p-value < 0.01) correlations between social behavior and stubb scores in gene promoters. We split this set of correlations into those with positive correlations with social species and those with positive correlations with solitary species. If social species have a greater capacity for gene regulation compared to lineages that have reverted to solitary nesting, then we would expect to find more motifs with scores that are higher in social taxa compared to secondarily solitary taxa. Only counting those motifs for which there were differences in numbers of significant correlations between social and solitary taxa of at least 20%, there were 94 motifs with more positive correlations with social taxa and 29 motifs with more positive correlations with solitary taxa. Five of the socially-biased motifs were previously associated with eusocial evolution in bees[7], including the motifs for *lola, hairy, CrebA, CG5180,* and the *met/tai* complex (Table S4).

This appears to be a large difference in the number of motifs positively correlated with social behavior and positively correlated with solitary behavior. We determined the significance of this result using a permutation test. We reran the PGLS analyses 100 times using six (of 15) randomly chosen species as focal (social) taxa. Only two of these permutations yielded at least 94 motifs with more positive correlations in these taxa. None yielded differences between the number of motifs with more positive correlations with social behavior and more positive correlations with solitary behavior as or more extreme than our result (94-29=65). Therefore, the pattern found appears to be significantly more extreme than expected by chance (Permutation test p-value < 0.01).

#### Greater constraint on noncoding elements in social species

Interestingly, genes showing relaxed selection associated with the loss of eusociality are enriched for chromosome condensation (Table S6), suggesting there may be trade-offs or costs associated with this increased regulatory potential. To determine whether or not we see a signature of these trade-offs in non-coding sequences, we used phastCons[82] to identify conserved non-exonic elements (CNEEs) in each species. We identified CNEEs that were either slow- or fast- evolving in eusocial and solitary lineages. Using the logic outlined above, we would predict that CNEEs essential to the maintenance of social behavior should be more highly conserved and evolving more slowly in extant eusocial taxa vs. secondarily solitary taxa. After phylogenetic correction, we find 1,876 CNEEs that meet these criteria, with faster rates of evolution in secondarily solitary species (Fig. 2c). This is significantly more than expected by chance (Binomial test, $p=5.27e^{-9}$); these regions are proximal to genes enriched for several GO terms, including serotonin receptor signaling (Permutation tests, p=0.007) and adherens junction maintenance (Permutation tests, p=0.009; see Table S15 for complete list). We also find 1,255 CNEEs that are more highly conserved in secondarily solitary lineages (fast in eusocial; slower in solitary). This number is smaller than expected by chance (Binomial test, $p<1e^{-10}$), but these regions are proximal to genes enriched for cytoskeletal organization and processes related to synapse and neurotransmitter functions, among other GO terms (Permutation tests, p<0.05; Table S15), potentially providing molecular support to hypotheses that the evolution of eusociality involves altered investments in cognitive function[141,142].

#### microRNA evolution

We identified a total of 1269 microRNAs (miRs) expressed in the brains of 15 species (including two populations of *L. albipes,* one solitary and one social) (Table S2). Most miRs were located in intergenic regions, and target predictions from Miranda and RNAhybrid identified gene targets for all but five sequenced miRs. Predicted targets of miRs were enriched for many GO terms (1857 terms for all miR targets; 1352 terms for targets in social species; 1148 terms for targets in solitary species). Social state of species did not predict the number of miRs expressed, nor the number of targets present across miRs (PGLS, p>0.05 in all comparisons).

There was weak but significant overlap for genes under positive selection at the origin of eusociality for Augochlorini and predicted miR targets in *Augochlorella aurata* (Hypergeometric test; RF=1.2, p=0.024), but not for other social species in Augochlorini. No significant overlap was found between genes under selection at the Halictini origin of eusociality and miR targets of any species. No significant overlap was found between genes under relaxation of selection associated with solitary reversions and miR targets of any species.

#### Development of LC/MS method for detection of JH III

We used LC-MS to measure JH III titers across tissues in *Bombus impatiens* as a proof of concept. JH III was detected in multiple bumblebee tissue types, including the brain, hemolymph, and ovaries (Fig. S6). Very low levels of JH (similar to dissection solution control) were detected in the retrocerebral (RCC) complex, which synthesizes JH (Fig. S6). The RCC immediately releases and is not known to store JH[143]. To determine whether we could track the movement of JH III through bee tissues, we applied JH III labeled with deuterium isotopes (JH III-D3) to the ventral abdominal sternites of *B. impatiens* workers. Labelled JH III-D3 levels were high in both bee bodies and the brain after 24 hours, but decayed significantly by 72 hours in both tissues (Fig. S7). JH III-D3 accounts for nearly all the total JH III in bee bodies after 24 hours, indicating that the labelled compound is well-absorbed by the bee (Fig. S7). Details of JH III and JH III-D3 quantification are provided in Fig. S8. Similar methods were used for detection of JH III and JH III-d3 in *A. aurata* and *A. pura,* results of which are presented in the main text.

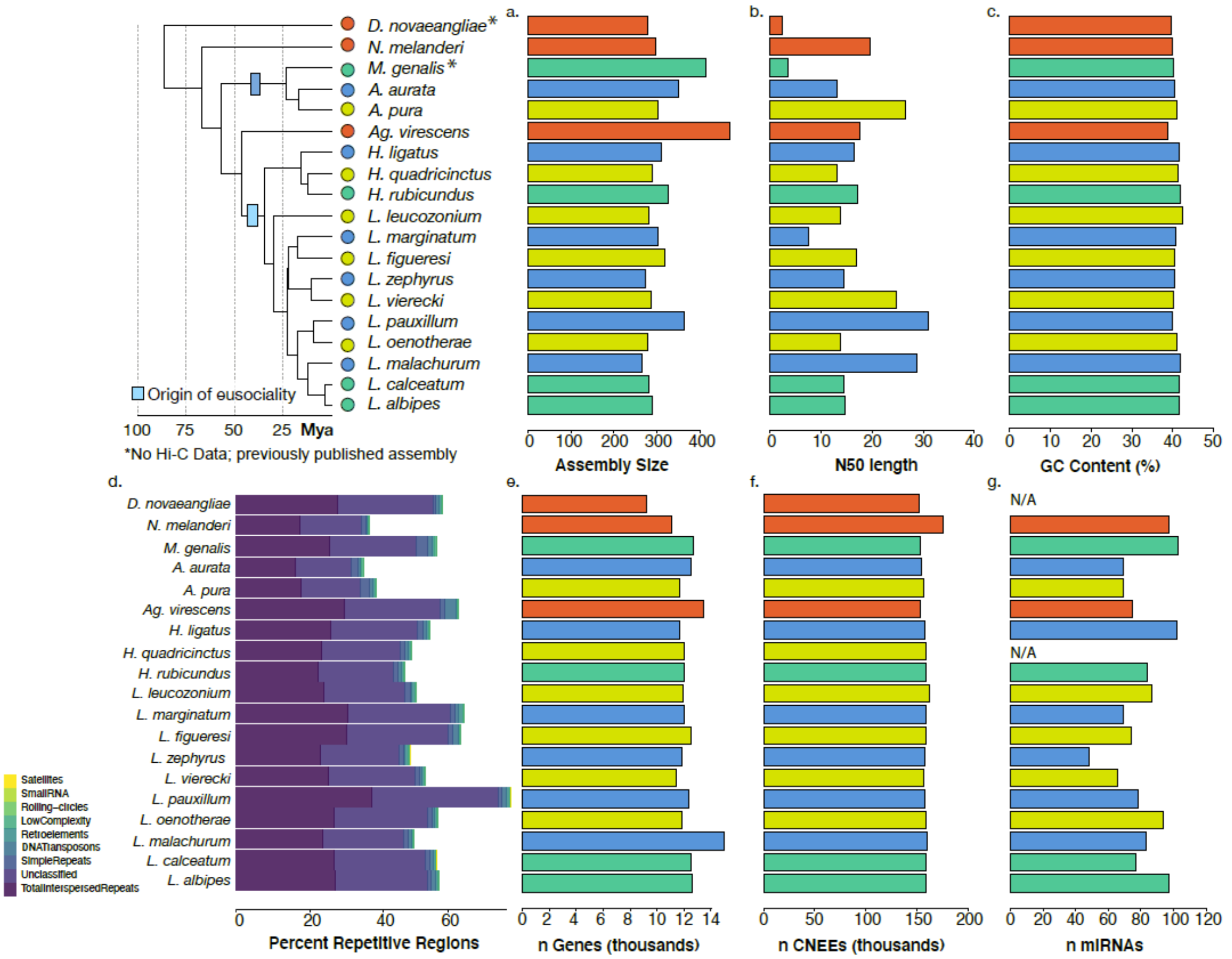


**Fig. S1.**

Genome assembly statistics. Nineteen genomes are included in this comparative dataset. *15 de novo* assemblies were generated using a combination of 10x genomics and/or Hi-C, and 2 previously-published genomes (*N. melanderi*[144], *L. albipes*[76]) were improved by scaffolding with Hi-C data. *M. genalis*[145] and *D. novaeangliae*[7] were used as-is. (**A**) Genome assembly lengths ranged from ~300-420Mb. (**B**) Scaffold N50s for each species following Hi-C scaffolding, and (**C**) GC content was consistent across species. (**D**) RepeatModeler was used to characterize different types of repeats in the halictid assemblies. (**E**) Numbers of genes (in thousands) for each species following individual annotation. (**F**) Conserved non-exonic elements (CNEEs) were called using phastCons on a progressive Cactus alignment. (**G**) microRNAs were also characterized using brain tissue from available species and from[92]. For some species, fresh tissue was not available (N/A).

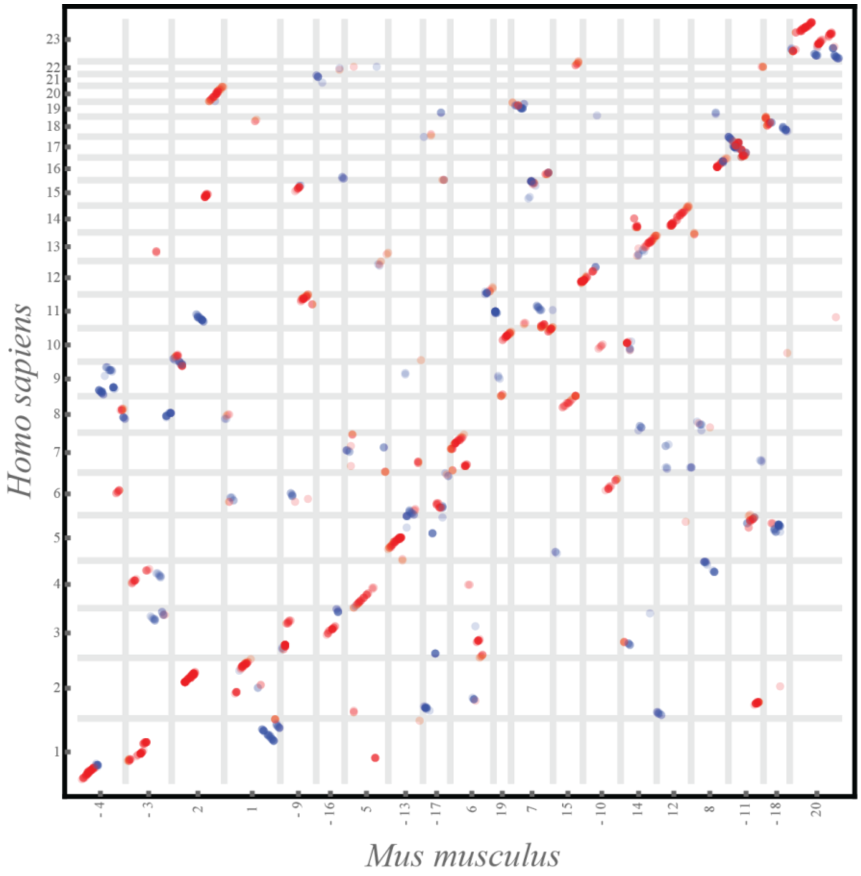


**Fig. S2.**

Dotplots comparing the chromosomes of two mammalian species, human and mouse, separated by comparable evolutionary distance to the bees examined in this study (~80MY to common ancestor). The vertical and horizontal lines outline the boundaries of chromosomes in respective species, and the numbers on the axes mark the relevant chromosome name and orientation, with “-“ in front of the chromosome name representing a reverse complement to the chromosome sequence as reported in the assembly. See Fig. S3 for more details. The human-mouse one-to-one alignments file was downloaded from https://github.com/mcfrith/last-genome-alignments.

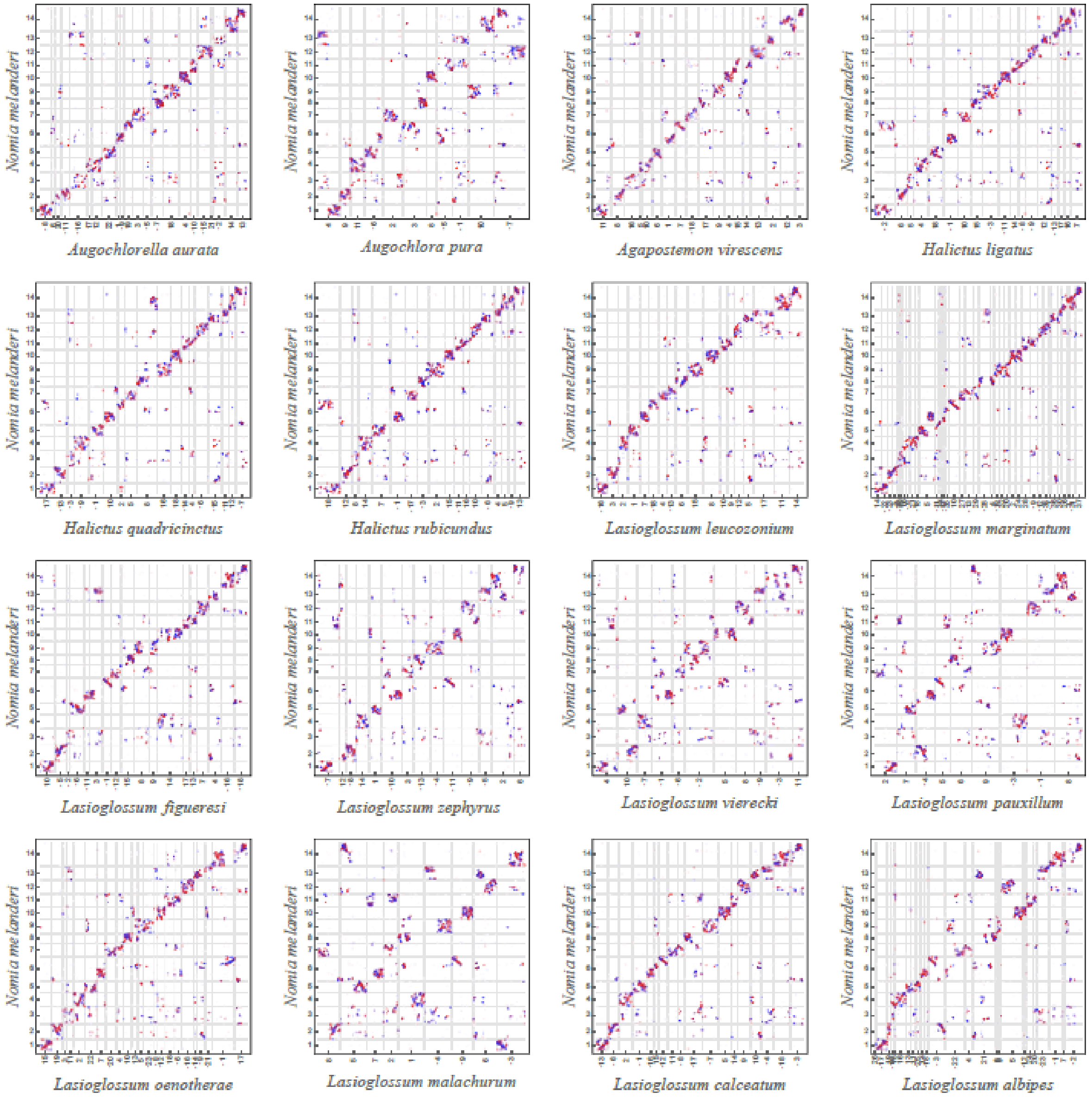


**Fig. S3.**

Dotplots showing alignment of chromosome-length scaffolds from 16 bee assemblies against the *N. melanderi* (NMEL) chromosome-length genome assembly. The NMEL reference (generated as part of this study) is shown on the y-axis. The x-axis shows the chromosome-length scaffolds in the respective bee assemblies that have been ordered and oriented to best match the NMEL chromosomes in order to facilitate comparison. The vertical and horizontal lines outline the boundaries of chromosomes in respective species, and the numbers on the axes mark the relevant chromosome name and orientation, with "-" in front of the chromosome name representing a reverse complement to the chromosome sequence as reported in the assembly. Each dot represents the position of a 1000 bp syntenic stretch between the two genomes identified by Progressive Cactus alignments. The color of the dots reflects the orientation of individual alignments with respect to NMEL (red indicates collinearity, whereas blue indicates inverted orientation). The dotplots illustrate that, with the exception of a few species, highly conserved regions belonging to the same chromosome in one species tend to lie on the same chromosome in other bee species, even though their relative position within a chromosome may change dramatically. This rearrangement pattern accounts for the characteristic appearance of the dotplots with large diffuse blocks of scrambled chromosome-to-chromosome alignments appearing along the diagonal. The pattern is visibly different from those characteristic of mammalian chromosome evolution (e.g., see Fig. S2). The few exceptions are species with multiple fissions (*L. marginatum*, *L. albipes*) and fusions (*Augochlora pura*, *L. vierecki*, *L. pauxillum*, *L. malachurum*) events. In the fission species, the syntenic regions that belonged to two different chromosomes in one bee species tend to belong to different chromosomes after the fission. The analysis of the fusion species suggests that the regions corresponding to separate chromosomes in the closely related species (and likely the ancestral species) remain separate in the new genome, possibly corresponding to the two chromosome arms. The alignments have been extracted from the hal file using the cactus hal2maf utility with the following parameters: --maxRefGap 500 --maxBlockLen 1000 --refGenome NMEL.

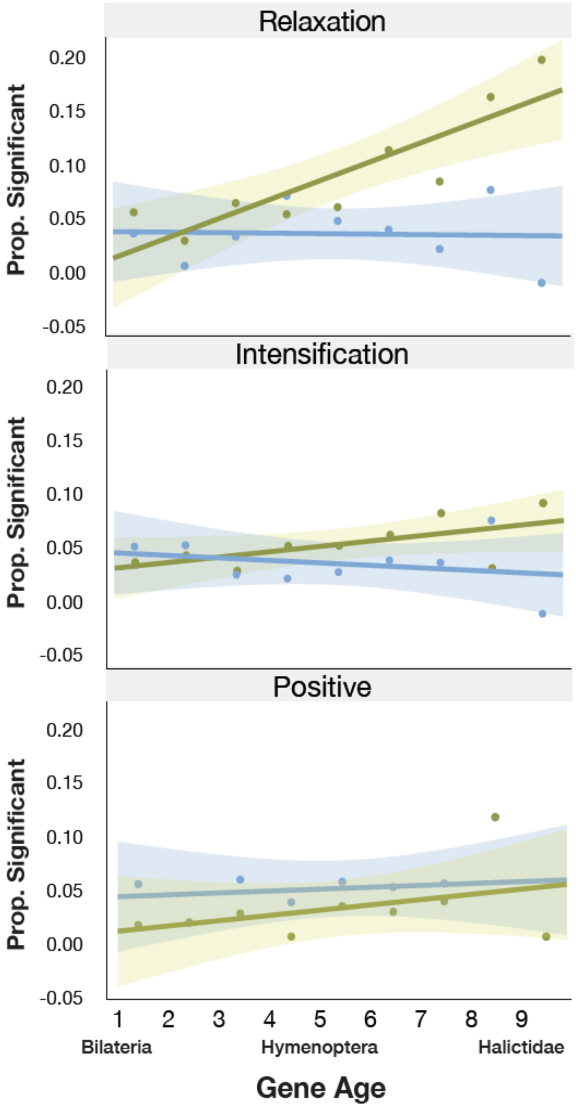


**Fig. S4.**

We examined the relationship between gene age and selection associated with eusocial origins, maintenance, and reversions to solitary life histories to determine whether or not there was any relationship between selection and gene age. Gene age ranges from the oldest Bilaterian group (Age=1) to the youngest, halictid-specific taxonomically restricted genes (Age=9). The relaxation panel demonstrates that there is a greater proportion of young genes experiencing relaxed selection when eusociality is lost (HyPhy RELAX, FDR<0.1; Pearson's r=0.869, p=0.002); we find no significant association with relaxation on extant, eusocial branches (Pearson correlation, r=-0.044, p=0.91). Next, we looked at the sets of genes that show intensification of selection pressures (HyPhy RELAX, FDR<0.1), but neither of these sets showed any significant association with gene age (Eusocial: Pearson correlation, r=0.244, p=0.492; Solitary: Pearson correlation, r=0.627, p=0.071). Finally, we looked at genes that showed evidence of positive selection (HyPhy aBSREL, FDR<0.05). We found no relationship between gene age and the proportion of genes with evidence of positive selection on at least 1 branch representing the origins of eusociality (Positive, gains: Pearson correlation, r=0.156, p=0.688). Likewise, there was no relationship between gene age and the proportion of genes with evidence for positive selection (HyPhy aBSREL, FDR<0.05) on at least 1 loss branch in the Halictini and on 1 loss branch in the Augochlorini (Positive, losses: Pearson correlation, r=0.403, p=0.283). Shading represents the 95% confidence intervals.

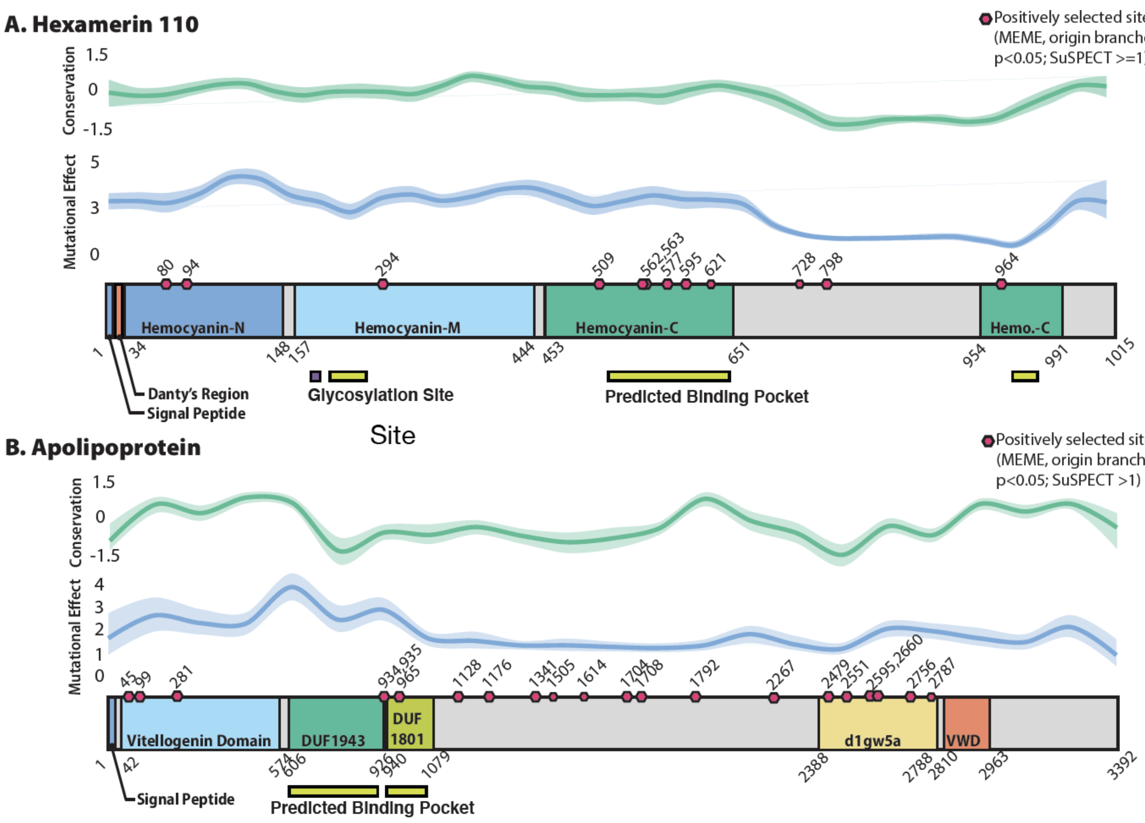


#### Fig. S5.

Both hex110 (**A**) and apolpp (**B**) show evidence for domain-specific positive selection associated with the origins of eusociality. Domain predictions were made with PHYRE[134,135]. Predicted binding pockets by PHYRE are shown with pink rectangles, glycosylation sites in orange squares. Branch-site specific tests for diversifying positive selection were conducted using the two branches associated with the origins of eusociality as focal branches (MEME[40], p<0.05). Pink hexagons denote MEME-identified sites that also had a mutational effect score (SuSpect[146]) >=1 for hex110 and >1 for apolpp. Conservation scores were estimated with AL2CO[147]. In both JHBPs, MEME identifies sites in functional regions of the protein, including the receptor binding domain and predicted binding pocket for ApoLpp as well as in all three Hemocyanin domains (associated with storage functions in these proteins) and in the predicted binding pocket of Hex110. Moreover, for both proteins, we also find region-specific, faster rates of evolution on eusocial branches compared to non-eusocial outgroups in this phylogeny (data not shown): the predicted binding pocket for ApoLpp and all Hemocyanin domains for Hex110. Taken together, these results suggest that positive selection shaped protein function as eusociality emerged in this group of bees.

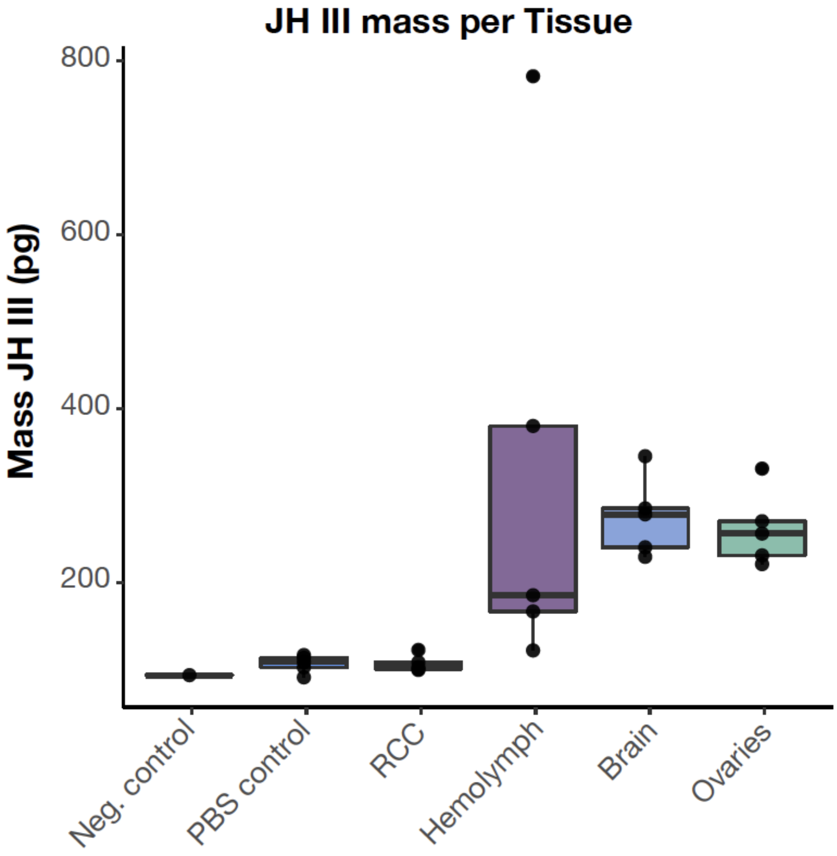


**Fig. S6.**

JH III is present in multiple bumblebee tissue types, including the brain. LC-MS was used to measure JH III levels in dissected brain tissue and hemolymph from worker bumblebees (Koppert). Tissues were dissected in PBS. Negative control=fresh PBS, PBS control=3uL of buffer collected following brain dissections from each sample. RCC=retrocerebral complex. The RCC synthesizes JH and immediately releases it. It is not known to store JH[143]. Hemolymph (3uL) was collected by centrifuging thorax tissue from each sample. Ovaries and brains were dissected in PBS and the entire organ was used for JH III quantification. In all samples, we find detectable levels of JH III in the hemolymph, brain, and ovaries. Note that because all samples were generated by extraction in a constant volume of buffer from the total input tissue, estimated amounts are not directly comparable across different tissue types.

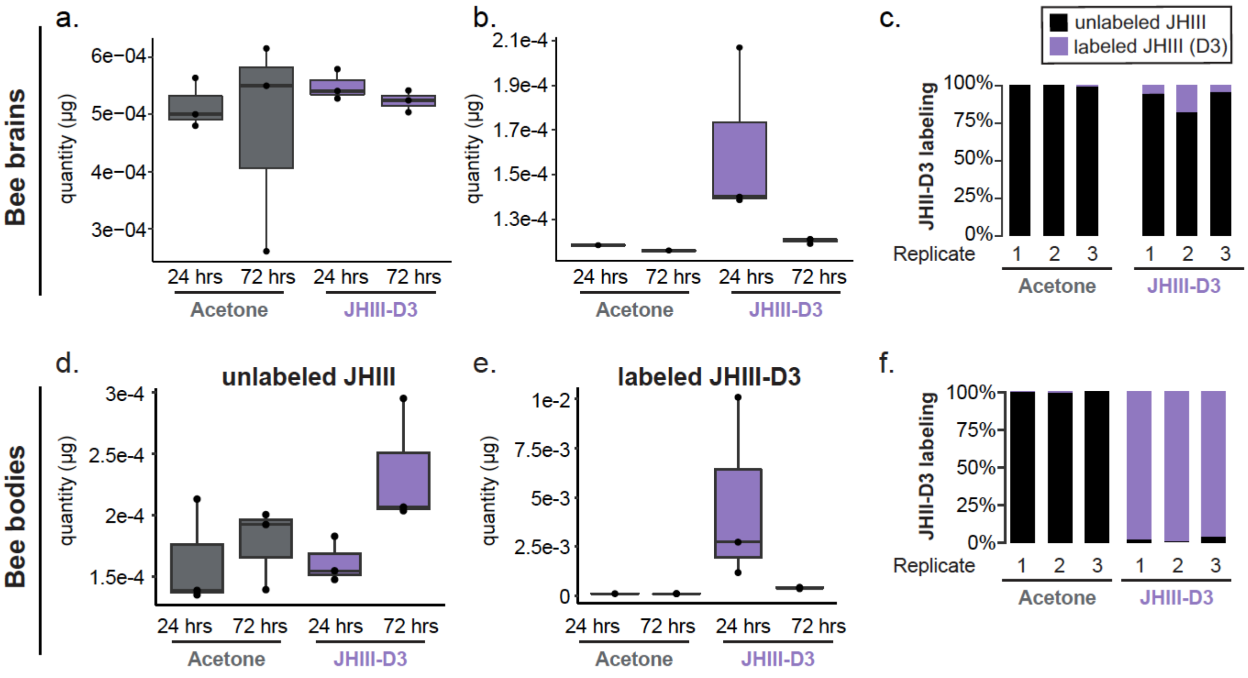


#### Fig. S7.

Stable Isotope Tracing Absolute quantification of JH III in the brain (**A**) and bodies (**D**) of bumblebees treated with acetone or JH III-D3. Absolute quantification of labelled JH III in the brains (**B**) and bodies (**E**) of bees treated with acetone or JH III-D3. Labelled JH III-D3 levels are high after 24 hours, but decay significantly by 72 hours. (**C**) Labelled JH III-D3 applied to abdomens of bumblebees is detectable in brains 24h later (acetone is control). n=3 tissues/condition. (**F**) JH III-D3 accounts for nearly all the total JH III in bee bodies after 24 hours, indicating that the labelled compound is well-absorbed by the bee. n=3 bumblebee workers for each experimental condition. Boxes show median and 25th and 75th percentiles. Whiskers show minimum and maximum values.

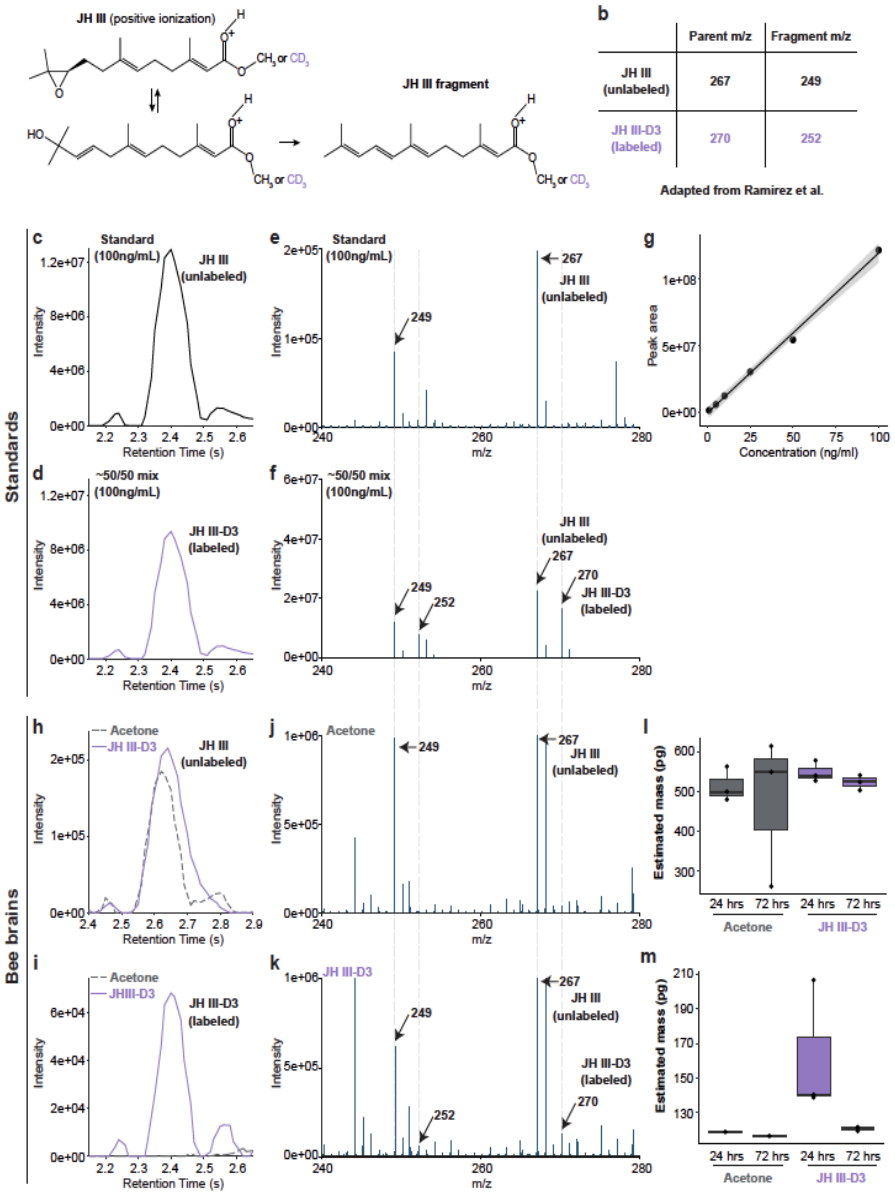


#### Fig. S8.

JH III and JH III-D3 quantification. **(A)** Chemical structure of JH III in positive ionization. JH III can exist in two forms of equal m/z and produces one fragment that retains the D3 label. **(B)** Parent and fragment m/z values for unlabelled and labelled JH III. **(C)** Chromatograph of unlabelled JH III. **(D)** Chromatograph of labelled JH III showing similar retention time to unlabelled JH III. **(E)** Mass spectra of unlabelled JH III showing the expected masses based on B. **(F)** Mass spectra of a 50/50 mix of labelled and unlabelled JH III showing the expected masses based on B. **(G)** Standard curve demonstrating a range of detection of JH III. **(H)** Chromatograph of unlabelled JH III in the brains of bees painted with acetone (grey dotted line) or with JH III-D3 (purple solid line). **(I)** Chromatograph of labelled JH III in the brains of bees painted with acetone (grey dotted line) or with JH III-D3 (purple solid line). **(J)** Mass spectra of JH III in bee brains painted with acetone showing the expected masses based on B. **(K)** Mass spectra of JH III in bee brains painted with JH III-D3 showing the expected masses based on B. **(L)** Absolute quantification of JH III in the brains of bees painted with acetone or JH III-D3. **(M)** Absolute quantification of JH III-D3 in the brains of bees painted with acetone or JH III-D3. n=3 replicates for each panel. Boxes show median and 25th and 75th percentiles. Whiskers show minimum and maximum values.

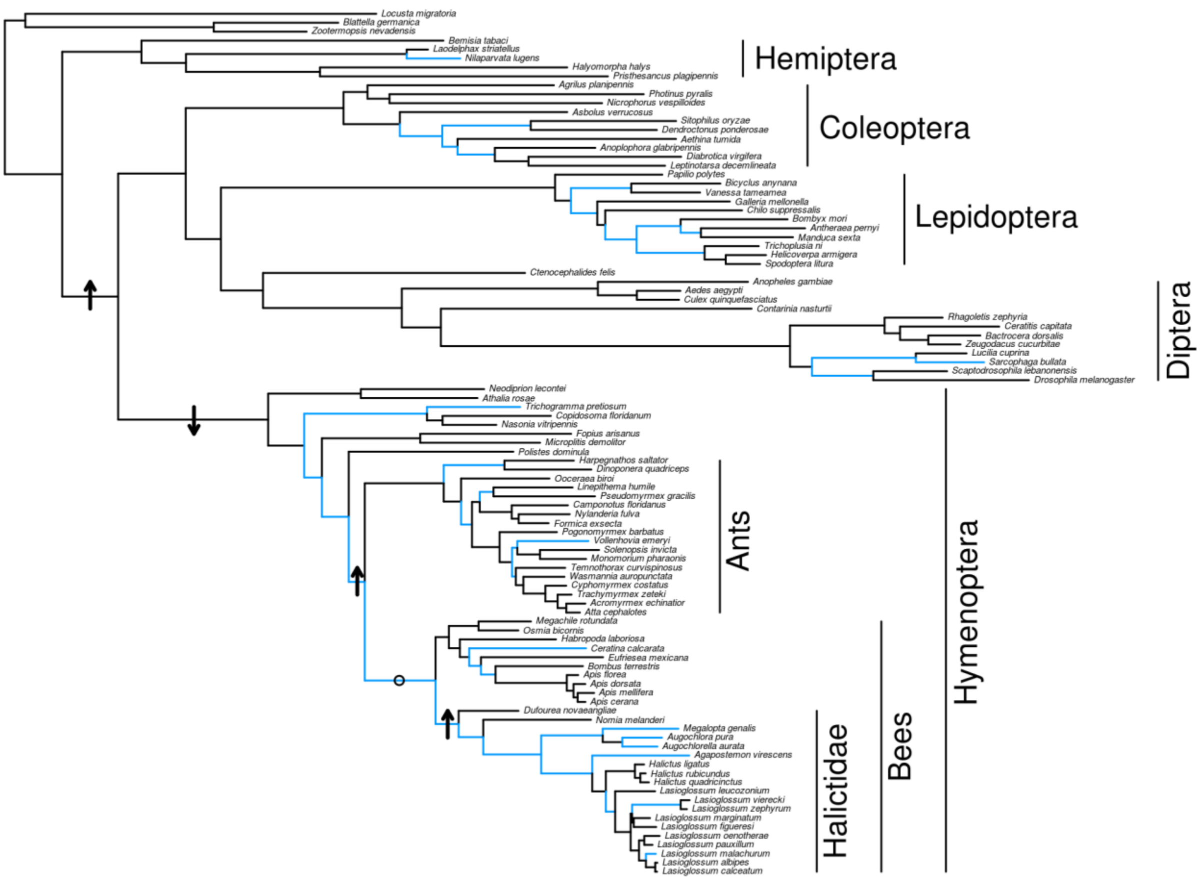


**Fig. S9.**

Apolipoprotein has experienced pervasive positive selection throughout the insects. 97 coding sequences were used to generate an insect-wide apolpp phylogeny (Table S11). HyPhy aBSREL was used to identify branches with evidence of positive selection (highlighted in blue). Branch-specific rate shifts in amino acid evolution were identified using likelihood ratio tests in PAML[86,87]; methods as in[148]. Arrows indicate the direction of significant rate shifts detected on relevant branches. The only branch tested that did now show a significant rate shift is indicated by an "o".

**Table S1. (separate file)**

Genome annotation statistics. Species name, total assembly length, karyotype, number of genes included in the assembly, the number of isoforms, and BUSCO scores.

**Table S2. (separate file)**

Characterization of microRNAs expressed in the brains of each species/population.

**Table S3. (separate file)**

Correlations between gene age and the proportion of gene sets of interest of each age.

**Table S4. (separate file)**

TF motif results in promoter regions of social vs. solitary lineages.

**Table S5. (separate file)**

Orthogroups with significant signatures of selection.

**Table S6. (separate file)**

GO enrichments among sets of coding sequences with signatures of selection. Parameter provides information on which test was conducted (RELAX or aBSREL), selection type denotes which branches were considered, and/or relevant significance thresholds.

**Table S7. (separate file)**

Samples included in this study. Sample name, species identification, collection date and location, sex of the individual, library preparation method, BioSample and BioProject accession IDs.

**Table S8. (separate file)**

Gene expression data for each species and tissue type.

**Table S9. (separate file)**

Cluster-specific DEGs (FDR < 0.05; 10 PCs, Resolution=0.6) for all 6716 cells in halictid scRNA-Seq experiment

**Table S10. (separate file)**

Cluster-specific DEGs (FDR < 0.05;6 PCs, Resolution=0.3) for all cells in Cluster 6 (glial cluster)

**Table S11. (separate file)**

Sequences included in evolutionary analyses for apolpp across insects.

**Table S12. (separate file)**

Hypergeometric tests examining enrichment of tested gene sets.

**Table S13. (separate file)**

Wilcoxon rank-sum tests of differences in tissue specificity between the genes of interest and all other genes.

**Table S14. (separate file)**

Numbers of branches under selection in apolpp across insects.

**Table S15. (separate file)**

GO enrichments among sets of CNEEs as determined by permutation tests.

#### Supplementary references.


60. Rao, S. S. P. *et al.* A 3D map of the human genome at kilobase resolution reveals principles of chromatin looping. *Cell* **159**, 1665–1680 (2014).
61. Davie, K. *et al.* A Single-Cell Transcriptome Atlas of the Aging Drosophila Brain. *Cell* **174**, 982-998.e20 (2018).
62. Lu, W., Wang, L., Chen, L., Hui, S. & Rabinowitz, J. D. Extraction and Quantitation of Nicotinamide Adenine Dinucleotide Redox Cofactors. *Antioxid. Redox Signal.* **28**, 167–179 (2018).
63. Wang, L. *et al.* Peak annotation and verification engine (PAVE) for untargeted LC-MS metabolomics. *Anal. Chem.* **91**, 1838–1846 (2019).
64. Martin, M. Cutadapt removes adapter sequences from high-throughput sequencing reads. *EMBnet.journal* **17**, 10–12 (2011).
65. Weisenfeld, N. I., Kumar, V., Shah, P., Church, D. M. & Jaffe, D. B. Direct determination of diploid genome sequences. *Genome Res.* **27**, 757–767 (2017).
66. Chakraborty, M., Baldwin-Brown, J. G., Long, A. D. & Emerson, J. J. Contiguous and accurate de novo assembly of metazoan genomes with modest long read coverage. *Nucleic Acids Res.* **44**, e147 (2016).
67. Waterhouse, R. M. *et al.* BUSCO Applications from Quality Assessments to Gene Prediction and Phylogenomics. *Mol. Biol. Evol.* **35**, 543–548 (2018).
68. Seppey, M., Manni, M. & Zdobnov, E. M. BUSCO: Assessing Genome Assembly and Annotation Completeness. *Methods Mol. Biol.* **1962**, 227–245 (2019).
69. Jackman, S. D. *et al.* ABySS 2.0: resource-efficient assembly of large genomes using a Bloom filter. *Genome Res.* **27**, 768–777 (2017).
70. Bioinformatics, B. FastQC: a quality control tool for high throughput sequence data. *Cambridge, UK: Babraham Institute* (2011).
71. Adey, A. *et al.* In vitro, long-range sequence information for de novo genome assembly via transposase contiguity. *Genome Res.* **24**, 2041–2049 (2014).
72. Zheng, G. X. Y. *et al.* Haplotyping germline and cancer genomes with high-throughput linked-read sequencing. *Nat. Biotechnol.* **34**, 303–311 (2016).
73. Burton, J. N. *et al.* Chromosome-scale scaffolding of de novo genome assemblies based on chromatin interactions. *Nat. Biotechnol.* **31**, 1119–1125 (2013).
74. Grabherr, M. G. *et al.* Full-length transcriptome assembly from RNA-Seq data without a reference genome. *Nat. Biotechnol.* **29**, 644–652 (2011).
75. Kim, D., Paggi, J. M., Park, C., Bennett, C. & Salzberg, S. L. Graph-based genome alignment and genotyping with HISAT2 and HISAT-genotype. *Nat. Biotechnol.* **37**, 907–915 (2019).
76. Kocher, S. D. *et al.* The draft genome of a socially polymorphic halictid bee, Lasioglossum albipes. *Genome Biol.* **14**, R142 (2013).
77. Smit, A. F. A. & Hubley, R. RepeatModeler Open-1.0. (2008).
78. Li, W. & Godzik, A. Cd-hit: a fast program for clustering and comparing large sets of protein or nucleotide sequences. *Bioinformatics* **22**, 1658–1659 (2006).
79. Paten, B. *et al.* Cactus: Algorithms for genome multiple sequence alignment. *Genome Res.* **21**, 1512–1528 (2011).
80. Sackton, T. B. *et al.* Convergent regulatory evolution and loss of flight in paleognathous birds. *Science* **364**, 74–78 (2019).
81. Hubisz, M. J., Pollard, K. S. & Siepel, A. PHAST and RPHAST: phylogenetic analysis with space/time models. *Brief. Bioinform.* **12**, 41–51 (2011).
82. Siepel, A. *et al.* Evolutionarily conserved elements in vertebrate, insect, worm, and yeast genomes. *Genome Res.* **15**, 1034–1050 (2005).
83. Earl, D. *et al.* Alignathon: a competitive assessment of whole-genome alignment methods. *Genome Res.* **24**, 2077–2089 (2014).
84. Bradley, R. K. *et al.* Fast statistical alignment. *PLoS Comput. Biol.* **5**, e1000392 (2009).
85. Capella-Gutiérrez, S., Silla-Martínez, J. M. & Gabaldón, T. trimAl: a tool for automated alignment trimming in large-scale phylogenetic analyses. *Bioinformatics* **25**, 1972–1973 (2009).
86. Yang, Z. PAML: a program package for phylogenetic analysis by maximum likelihood. *Comput. Appl. Biosci.* **13**, 555–556 (1997).
87. Yang, Z. PAML 4: phylogenetic analysis by maximum likelihood. *Mol. Biol. Evol.* **24**, 1586–1591 (2007).

115. Dohrmann, M. & Wörheide, G. Dating early animal evolution using phylogenomic data. *Sci. Rep.* **7**, 3599 (2017).
116. Patro, R., Duggal, G., Love, M. I., Irizarry, R. A. & Kingsford, C. Salmon provides fast and bias-aware quantification of transcript expression. *Nat. Methods* **14**, 417–419 (2017).
117. Bolstad, B. M. & Bolstad, M. B. M. Package 'preprocessCore.' (2013).
118. Pennell, M. W. *et al.* geiger v2.0: an expanded suite of methods for fitting macroevolutionary models to phylogenetic trees. *Bioinformatics* **30**, 2216–2218 (2014).
119. Chen, S., Zhou, Y., Chen, Y. & Gu, J. fastp: an ultra-fast all-in-one FASTQ preprocessor. *Bioinformatics* **34**, i884–i890 (2018).
120. Hao, Y. *et al.* Integrated analysis of multimodal single-cell data. *Cell* **184**, 3573-3587.e29 (2021).
121. Hafemeister, C. & Satija, R. Normalization and variance stabilization of single-cell RNA-seq data using regularized negative binomial regression. *Genome Biol.* **20**, 296 (2019).
122. Finak, G. *et al.* MAST: a flexible statistical framework for assessing transcriptional changes and characterizing heterogeneity in single-cell RNA sequencing data. *Genome Biol.* **16**, 278 (2015).
123. Traniello, I. M. *et al.* Meta-analysis of honey bee neurogenomic response links Deformed wing virus type A to precocious behavioral maturation. *Sci. Rep.* **10**, 3101 (2020).
124. Sheng, L. *et al.* Social reprogramming in ants induces longevity-associated glia remodeling. *Sci Adv* **6**, eaba9869 (2020).
125. Yin, J. *et al.* Brain-specific lipoprotein receptors interact with astrocyte derived apolipoprotein and mediate neuron-glia lipid shuttling. *Nat. Commun.* **12**, 2408 (2021).
126. Fisher, C. E. & Howie, S. E. M. The role of megalin (LRP-2/Gp330) during development. *Dev. Biol.* **296**, 279–297 (2006).
127. DeSalvo, M. K. *et al.* The Drosophila surface glia transcriptome: evolutionary conserved blood-brain barrier processes. *Front. Neurosci.* **8**, 346 (2014).
128. Melom, J. E. & Littleton, J. T. Mutation of a NCKX eliminates glial microdomain calcium oscillations and enhances seizure susceptibility. *J. Neurosci.* **33**, 1169–1178 (2013).
129. Stork, T., Bernardos, R. & Freeman, M. R. Analysis of glial cell development and function in Drosophila. *Cold Spring Harb. Protoc.* **2012**, 1–17 (2012).
130. Zhang, H. *et al.* Drosophila CG10527 mutants are resistant to juvenile hormone and its analog methoprene. *Biochem. Biophys. Res. Commun.* **401**, 182–187 (2010).
131. Yanai, I. *et al.* Genome-wide midrange transcription profiles reveal expression level relationships in human tissue specification. *Bioinformatics* **21**, 650–659 (2005).
132. Jarvis, E. D. *et al.* Whole-genome analyses resolve early branches in the tree of life of modern birds. *Science* **346**, 1320–1331 (2014).
133. Le Guilloux, V., Schmidtke, P. & Tuffery, P. Fpocket: an open source platform for ligand pocket detection. *BMC Bioinformatics* **10**, 168 (2009).
134. Kelley, L. A., Mezulis, S., Yates, C. M., Wass, M. N. & Sternberg, M. J. E. The Phyre2 web portal for protein modeling, prediction and analysis. *Nat. Protoc.* **10**, 845–858 (2015).
135. Kelley, L. A. & Sternberg, M. J. E. Protein structure prediction on the Web: a case study using the Phyre server. *Nat. Protoc.* **4**, 363–371 (2009).
136. Edgar, R. C. MUSCLE: a multiple sequence alignment method with reduced time and space complexity. *BMC Bioinformatics* **5**, 113 (2004).
137. Edgar, R. C. MUSCLE: multiple sequence alignment with high accuracy and high throughput. *Nucleic Acids Res.* **32**, 1792–1797 (2004).
138. Pahlke, S., Seid, M. A., Jaumann, S. & Smith, A. The Loss of Sociality Is Accompanied by Reduced Neural Investment in Mushroom Body Volume in the Sweat Bee Augochlora Pura (Hymenoptera: Halictidae). *Ann. Entomol. Soc. Am.* **114**, 637–642 (2021).
139. Shpigler, H. Y. *et al.* Deep evolutionary conservation of autism-related genes. *Proc. Natl. Acad. Sci. U. S. A.* **114**, 9653–9658 (2017).
140. Roth, Z. *et al.* Identification of receptor-interacting regions of vitellogenin within evolutionarily conserved β-sheet structures by using a peptide array. *Chembiochem* **14**, 1116–1122 (2013).
141. Dunbar, R. I. M. The social brain hypothesis. *Evol. Anthropol.* **6**, 178–190 (1998).
142. O'donnell, S. *et al.* Distributed cognition and social brains: reductions in mushroom body investment accompanied the origins of sociality in wasps (Hymenoptera: Vespidae). *Proceedings of the Royal Society B: Biological Sciences* **282**, 20150791 (2015).
143. Frederik Nijhout, H. *Insect Hormones*. (Princeton University Press, 1998).
